# Hierarchy in Neuronal Representations of Multiple Tasks in Prefrontal Cortex

**DOI:** 10.64898/2026.02.10.705211

**Authors:** Qing Sheng, Sizheng Luo, Da Li, Jing Jia, Zixuan Fan, Zhenliang He, Fei Wang, Yulei Chen, Shuye Yuan, Zhixian Cheng, Chengyu T. Li, Yang Xie

**Affiliations:** Lingang Laboratory; Shanghai, 201101, China; Institute of Neuroscience, CAS Center for Excellence in Brain Science and Intelligence Technology, Chinese Academy of Sciences; Shanghai, 200031, China; Shenzhen Medical Academy of Research and Translation (SMART); Shenzhen, 518107, China; Global Institute of Future Technology, Shanghai Jiao Tong University; Shanghai, 200240, China

## Abstract

The ability to perform multiple tasks is one of the fundamental hallmarks of general intelligence in the brain. However, the underlying neural mechanisms remain largely unknown. We trained macaque monkeys to perform four tasks with identical spatial layouts but different cognitive demands, and recorded activities of thousands of prefrontal neurons using two-photon calcium imaging. Within the same neuronal population, we identified a multitask neural geometry composed of multiple low-dimensional subtask spaces, each encoding spatial information within a specific subtask. These subtask spaces shared a ring-like representational structure, forming a generalized spatial code across different subtasks. Task separation arose from separable bases and offsets of subtask spaces, modulated primarily by meta-task features. Thus, the neural geometry is organized by a three-level hierarchical structure: location codes are nested within subtask spaces, which are grouped as different meta-tasks. This hierarchy supported generalization across both locations and tasks, and explained monkeys’ error behaviors. Together, the hierarchically organized neural geometry underlies flexible multitask behavior.

## Introduction

Flexibly and effectively performing a wide variety of tasks is a defining feature of cognitive versatility and a cornerstone of general intelligence (*1*). While various machine learning methods have shown robust performance on specialized sets of multiple tasks (*2-5*), they typically require extensive training and still fall short of the broad, adaptive intelligence observed in biological systems (*1*). The brain, by contrast, exemplifies general intelligence through its remarkable ability to operate across diverse domains and goals. How does the brain represent and coordinate multiple tasks to achieve different goals?

Representation sharing reuses a compact set of underlying features or latent codes to support many tasks. Therefore, it can reduce redundant computation and memory storage. Both biological and artificial intelligence trained to perform multiple tasks will reuse representations and computational components across tasks (subtasks), exhibit generalizability across different tasks(*4, 6-10*), therefore potentially facilitating faster learning and inference for novel tasks(*11, 12*).

Crucially, however, widespread similarity across tasks can engender inter-task interference, degrading performance if all tasks compete for the same representational resources. This tension motivates the parallel importance of separation mechanisms. On single neuron level, distinct neuronal codes of task sets(*13*), each of which abstractly identifies individual task rules regardless of sensory features(*14, 15*), help to reduce the interference among different tasks. On population level, mutual interference of simultaneously memorized contents in working memory can be minimized by projecting relevant dimensions into orthogonal, low-dimensional subspaces(*16*). Such subspace organization permits parallel processing of multiple tasks while constraining interference.

To effectively and flexibly accomplish multiple tasks, the brain must exhibit both shared representations for high cross-task generalization and task-specific separation to minimize inter-task interference. However, these two features pull in opposite direction with same group of neurons. How the brain balanced both sharing and separation across multiple tasks is largely unknown.

The lateral prefrontal cortex (LPFC) plays a critical role in various cognitive tasks (*17-24*). Despite substantial progress in understanding the complex neural computations within the LPFC (*25-29*), it remains unclear how a single group of neurons in this region can represent multiple tasks. Specifically, the mechanisms by which these neurons achieve both representational sharing and representational separation are not well understood. Recent findings suggesting that cognitive processes are reflected in low-dimensional neural manifolds (*23, 30-33*) indicate that studying the geometry of multitask representations in the LPFC may provide valuable insights into the balance between shared and separate representations.

To investigate how multiple tasks are represented in the brain, we designed a multitask paradigm in which each task was comprised of a series of subtasks requiring monkeys to make decisions based on a fixed set of spatial locations, but within varying contexts involving go/no-go choices, immediate/delayed responses, attentional demands, reward-related memory, and inter-task interference. Using two-photon imaging, we recorded the activity of the same LPFC neurons across all tasks in two monkeys to address the following questions: 1) How do individual neurons represent multiple tasks? 2) What is the geometric representation of multiple tasks at the population level? 3) Whether and how does this neuronal geometry support both representation sharing and separation?

## Results

### Paradigm and behavior

Two monkeys (*Macaca fascicularis*) were trained to perform 4 tasks: a discrimination task, an attention task, a sequential reach task, and a nested reach task (Fig. 1a). These tasks required monkeys’ decisions at eight spatial locations under varying cognitive contexts, including go/no-go discrimination, congruent/incongruent spatial attention, reward memory, and inter-task interference. Each task was concatenated by series of subtasks. Collectively, the 4 tasks comprised a total of 14 subtasks. Both monkeys achieved high accuracies across all locations for every task, with average accuracies exceeding 0.75 (Fig. 1b). Detail descriptions of each task and the behavioral performance are provided below.

**Fig. 1.**
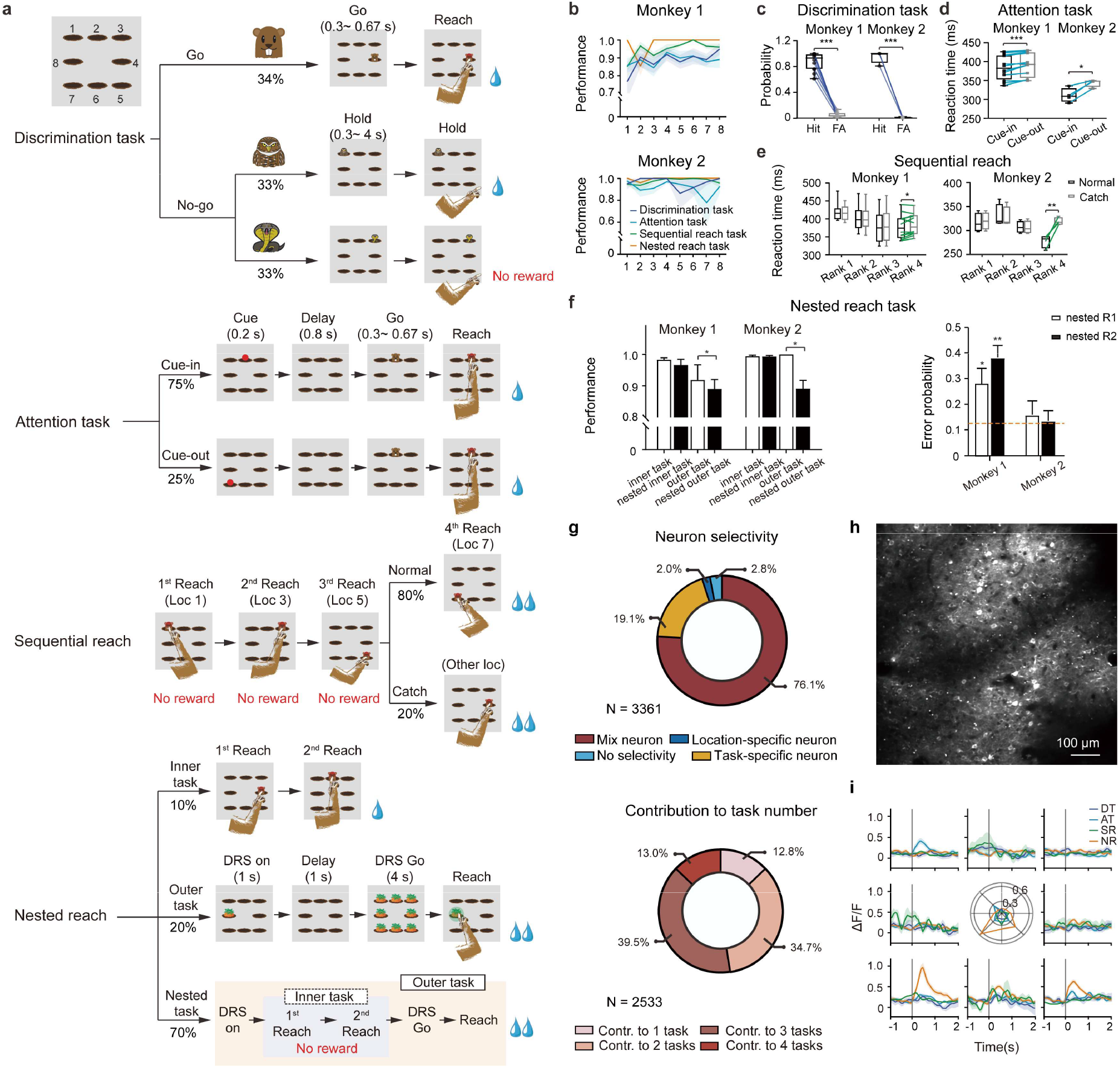
Task paradigm, behavior and single neuron activities. (**a**) Task structures (Methods). Loc, location; DRS, delay reach stimulus. The percentages indicate the proportions of different trial types. (**b**) Correct rates across tasks and locations for two monkeys. Each line represents a different task, with shaded areas indicating the standard error of the mean (SEM). Data were pooled across imaging sessions. (**c**) Discrimination task performance. Comparison of hit rates and FA rates for the discrimination task. Both monkeys showed significant differences between hit and FA rates (p values < 0.001, two-tailed paired t-test). Blue lines represent individual imaging sessions. (**d**) Attention task performance comparing RTs for cue-in and cue-out trials. Both monkeys exhibited significantly faster RTs in cue-in trials. (monkey 1: p < 0.001; monkey 2: p = 0.014, paired t-test). Light blue lines represent individual imaging sessions. (**e**) Sequential reach task performance comparing RTs for each reach in normal and catch trials. While the first three reaches showed no significant difference between trial types, the fourth reach revealed a significant difference. (monkey 1: p < 0.05; monkey 2: p < 0.01, two-tailed paired t-test). Data were averaged across imaging sessions. (**f**) Comparison of performance in inner and outer task trials, with and without nesting in nested reach task. Left: Correct rates for inner task, nested inner task, outer task, and nested outer task trials. Correct rates for the nested outer task were significantly lower than for the standalone outer task (monkey 1, 0.89 ± 0.03 vs. 0.92 ± 0.05, p = 0.03; monkey 2, 0.89 ± 0.03 vs. 1.0 ± 0, p = 0.03, Wilcoxon rank sum test). Right: Error responses in the nested reach task, showing the proportion of incorrect delay responses influenced by the cues for the first (NR_R1) and second (NR_R2) reaches. The orange line represents chance level (1/8). For all panels, error bars indicate SEM, *p < 0.05, **p < 0.01. (**g**) Neuron selectivity. Top: Distribution of selectivity types in LPFC neurons. Permutation tests classified LPFC neuron selectivity into four categories: no selectivity (2.8%), location-specific (2.0%), task-specific (19.1%), and mixed selectivity (76.1%). Bottom: Multitasking properties of mixed-selective neurons. Among mixed-selective neurons, 12.8% contributed to only one task, 34.7% contributed to two tasks, 39.5% contributed to three tasks, and 13.0% contributed to four tasks. (**h**) Illustration of two-photon calcium imaging of monkey LPFCs. AS: arcuate sulcus; PS: principal sulcus. (**i**) Example of a neuron with spatial selectivity, contributing to multiple tasks. Radar charts display the mean ΔF/F during the 0-670 ms period following go stimulus onset for each task (DT_Go: dark blue; AT_Go_in: light blue; SR_Go: green; NR_Go: orange). The radar chart directions correspond to the locations of 8 targets. Surrounding line charts show averaged neuronal responses for each location within each task, using matching colors. Shaded regions indicate SEM. The gray line indicates the onset of go stimulus. The x-axis represents time, and the y-axis represents ΔF/F.

In the discrimination task (DT), monkeys were required to categorize three visual stimuli into go and no-go groups. The go stimuli, represented by a mole image, required a touch to a specified screen location, while no-go stimuli required withholding a touch to the location indicated by owl or snake images. Correct responses to the go stimuli (hits) and correct no-go responses (correct rejections) to the owl image were rewarded with juice. Correct rejections to the snake image received no reward. Trials with these three types of stimuli were presented in random order. Behavioral performance was evaluated by comparing hit and false alarm (FA) rates for the go and no-go conditions. Both monkeys showed significantly higher hit rates than FA rates (Fig. 1c; permutation test, p < 0.001 for both monkeys), indicating successful go/no-go discrimination.

The attention task (AT) manipulated the monkeys’ spatial attention. Each trial began with a 200 ms attention cue, followed by an 800 ms interval before the reach target appeared. Monkeys were required to touch the target within 300-670 ms. In 75% of trials, known as cue-in trials, the attention and reach cues appeared on the congruent location. In the remaining cue-out trials, the attention and the reach cues appeared at incongruent locations. Reaction times (RTs) were significantly shorter for cue-in trials (monkey 1: 383.6 ± 30.6 ms; monkey 2: 310.5 ± 16.3 ms) compared to cue-out trials (monkey 1: 391.7 ± 29.3 ms; monkey 2: 343.6 ± 8.8 ms) in both monkeys (Fig. 1d; two-tailed paired t-test, monkey 1: p <0.001; monkey 2: p = 0.014), suggesting that the attention cue effectively modulated motor responses.

In the sequential reach task (SR), monkeys performed a fixed sequence of 4 reaches (locations 1-3-5-7) in 80% of trials. In the remaining catch trials, the final target was randomly chosen from the other seven locations excluding location 7 (locations 1-3-5-X). The order of locations allowed the monkeys to predict the remaining delay to reward availability. RTs for the fourth reach in catch trials were significantly longer than those in normal trials (Fig. 1e; two-tailed paired t-test, monkey 1: p < 0.05, monkey 2: p < 0.001), suggesting that the monkeys learned the structure of the standard sequence and anticipated the final location, thus leading to faster responses in normal trials.

The nested reach task (NR) embedded a double-reach task within the delay period of the delay reach stimulus (DRS) task. The outer task involved a 500 ms cue, followed by a 1000-2000 ms delay, during which monkeys had to touch the DRS cue’s location within 4000 ms after the go signal. The inner task required sequentially touching two reach cues within variable response windows (Extended Data Table 1). The NR task consisted of 10% standalone inner task trials, 20% standalone outer task trials, and 70% nested tasks trials, all randomly interleaved. Although overall performance on the NR task was good (Fig. 1f), subtle inter-task interference was observed. Accuracies on the outer task during the nested condition (monkey 1: 0.89 ± 0.03; monkey 2: 0.89 ± 0.03) were significantly lower than that during the standalone condition (monkey 1: 0.92 ± 0.05, monkey 2: 1.0 ± 0.0, p < 0.05 for both monkeys; Fig. 1f, left panel). Only monkey 2 exhibited significantly longer RTs in the nested condition (Extended Data Fig. 1a). For monkey 1, over 60% of incorrect delayed reaches in the nested task targeted the inner task locations (Fig. 1f, right panel, nested R1: p = 0.033, nested R2: p = 0.001, Wilcoxon signed-rank test). Although monkey 2’s errors were not significantly located on R1 or R2, they occurred around these locations more frequently than chance levels (Extended Data Figs. 1b-c). These results suggest the presence of inter-task interference during the nested reach task.

In summary, monkeys exhibited proficiency across distinct cognitive tasks, achieving high accuracy at all spatial locations. However, their performance varied depending on the task, with distinct patterns of accuracy and RTs at the same spatial locations. These behavioral findings suggest an interplay between shared spatial representations and context-dependent processing, highlighting the monkeys’ ability to implement specific rules within a unified spatial layout.

### Limitations of single neuron activities in capturing task sharing and separation

Is there any similarity in neural representation of spatial locations across different tasks? Can task separation at the behavior level also occur at the single-neuron level? One hypothesis posits that stable location representations are encoded by location-specific neurons, while task separation is supported by neurons specifically responding to task contexts. To assess this, we introduced the AAV virus expressing GCaMP6f into the LPFCs of two monkeys for two-photon imaging (Extended Data Figs. 1d-f; monkey 1: 2424 neurons from 12 field of views (FOVs), monkey 2: 937 neurons from 4 FOVs, Extended Data Fig. 1g).

To investigate the neural representation of identical spatial locations across 4 tasks, we selected one go stimulus onset event from each task: DT_Go, AT_Go_in, SR_Go, NR_Go (Methods). The neural responses averaged across time within each trial were analyzed. We found that only a small fraction of neurons (2.0%, Fig. 1j) specifically represented spatial locations. Most neurons exhibited mixed encoding of spatial locations and task contexts (76.1 %, Fig. 1j; Extended Data Figs. 1h-1i, example neurons). We further explored whether mixed-selective neurons were organized into distinct, task-dedicated groups. ANOVA results revealed that only a small proportion of mixed-selective neurons (12.8 %) were specialized for a single task (Extended Data Fig. 1h, example neuron), while the majority (87.2 %) contributed to multiple tasks (Fig. 1g; Fig. 1i, example multitasking neuron). Furthermore, the spatial tuning of neurons varied across tasks, showing not only changes in the scale of spatial tuning (Extended Data Fig. 1i) but also shifts in location preference (Fig. 1i).

Thus, these findings highlight the difficulty in isolating task-specific representations at the single-neuron level. The presence of mixed task representations in individual neurons is insufficient to account for the observed behavioral separation among tasks.

### The neural geometry of multiple tasks

We next examined how multiple tasks were organized at the population level. Each task in the multitask behavior comprised a varying number of subtasks. Thus, the neural representation of multiple tasks could be characterized by the geometric organization of neural states associated with spatial locations within the relevant subtasks. We hypothesized that the neural representation of spatial locations for each subtask was embedded in a specific low-dimensional subspace, which were defined as subtask spaces. And the neural evidence for task sharing and separation at the behavior level could be explained by studying these subtask spaces.

For each subtask, we first performed linear regression on the averaged neural responses during 330-1000 ms after the onset of related task events in correct trials. The linear regression model incorporated 8 spatial locations (encoded in a one-hot manner) and a bias term (Fig. 2a, Extended Data Eq. 1). For each subtask, we obtained vector representations of the population states for 8 locations by concatenating the regression coefficients of all neurons from both monkeys. We then applied principal component analysis (PCA) to the vectors within each subtask to extract a low-dimensional space (Extended Data Fig. 2a) capturing the major variance related to spatial locations (Methods).

**Fig. 2.**
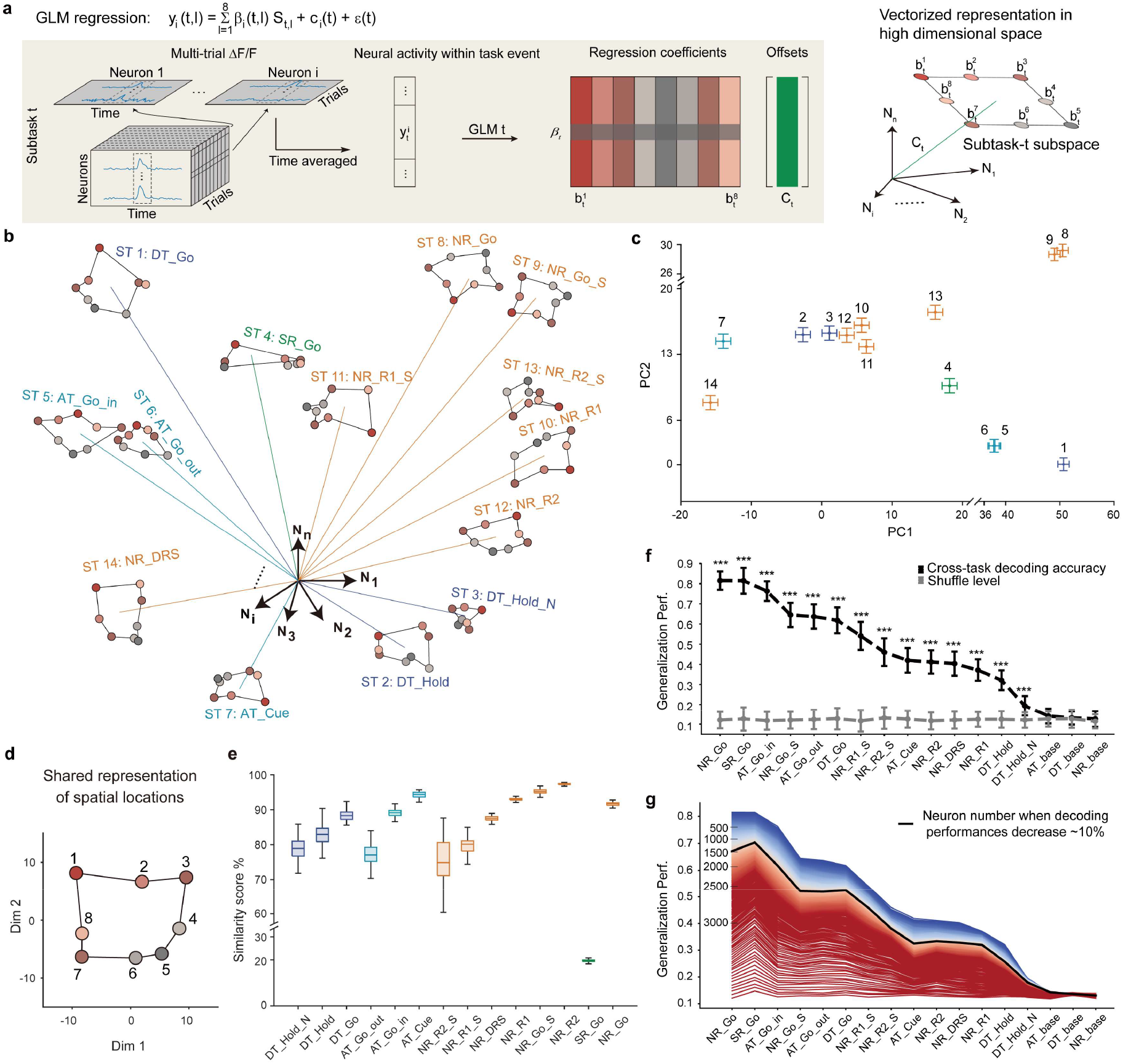
Geometric representation of multiple tasks. (**a**) Illustration of linear regressions on neural pseudo-populations for each subtask. Left: GLM analysis of neural responses for spatial locations within a subtask, using data from both monkeys. Right: Vectorized neural state representation. *N*_i_ represents the single neuron vector.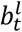, is the neural state for location *l* in subtask *t*, and *C*_t_ is the vectorized representation of the subtask-*t* space’s offset. (**b**) Population responses projected to the corresponding subtask space. Neural responses for task-location combinations, grouped after stimulus onset, were projected and color-coded by locations and subtasks. (**c**) Offsets of regression models from different subtasks. Offsets were projected onto a manifold characterizing subtask separation. PCA was applied to all neuron offsets. The number of subtasks is same with **(b)**, the error bars are mean ± 6 × SEM. (**d**) The shared representation of spatial locations was calculated from 12 subtasks (SR_Go and NR_Go were left out as tests). (**e**) Similarity scores of representational geometries of spatial locations across 14 subtasks. Colors indicate different subtasks, and error bars represent the standard deviation across 100 bootstraps. (**f**) Generalization performance on location discrimination across subtasks. A decoder, trained on data from SR_Go, NR_Go, AT_Go_in, and DT_Go, was tested on 14 subtasks and 3 baseline epochs. The gray curve indicates the level of trial shuffling. *p < 0.00294, **p < 0.00058, ***p < 0.00005 (permutation test with Bonferroni correction). All accuracies are presented as mean ± std. Error bars represent std across 100 bootstraps. (**g**) Results of incrementally shuffling neural signals in cross-subtask generalization tests. Neurons were shuffled in descending order based on their contribution to the decoder in (**f**). Line colors indicate an increasing shuffled neurons number. The black curve shows that shuffling the weights of the top 1500 contributing neurons leads to an approximately 10% decrease in generalization performance.

In addition, the offset of each subtask space was vectorized by concatenating the bias terms from the linear regression models of all neurons (denoted by *C*_*t*_ for subtask *t* in Fig. 2a). This allowed us to test our hypothesis by quantifying representational sharing through comparing the geometry of spatial location representations within each subtask space, and measuring the degree of separation among subtask spaces based on their orthonormal bases and offsets.

To investigate the neural geometry of multiple tasks, 14 two-dimensional subtask spaces were obtained in the high-dimensional neural state space (Fig. 2b and Extended Data Fig. 2b; Extended Data Fig. 2c, cumulative explained variance). In most subtask spaces, the neural states corresponding to different spatial locations exhibited a ring-like geometric structure. In addition to spatial locations, separable representations were also observed in the low-dimensional subtask-specific manifold (Fig. 2c), which captured the major variance of these offsets (Extended Data Fig. 2d, cumulative explained variance). This suggests that subtask-specific information can be characterized by the offsets of subtask spaces. This neural geometry also showed strong behavioral relevance, including incorrect go/no-go discrimination, attention modulation and inter-task interference (Extended Data Figs. 3).

As a result, we identified valid spatial codes embedded in each subtask space, characterized by distinct offsets within the multitask geometry.. Given the monkeys’ high accuracies across all locations in different tasks, both shared spatial codes and separate task codes were necessary. We then sought to assess both aspects of the geometric representation underlying multiple tasks.

### Representational sharing of spatial locations within the neural geometry

Effective task execution required the use of spatial information, so we investigated how the spatial locations were represented and whether these representations were shared across different subtasks. We found that neural representations in most subtask spaces exhibited similar ring-like structures (Fig. 2b and Extended Data Fig. 2b). Based on this, we hypothesized that the neural codes for spatial locations across subtask spaces were similar. By applying distinct affine transformations across 12 subtasks (excluding NR_Go and SR_Go as test sets, Extended Data Fig. 4a), we observed a shared representation structure (Fig. 2d, Methods), which mirrored the spatial layout presented to the monkeys. The similarity of spatial codes within each subtask was notably high, except for the SR_Go space, which was strongly influenced by reward memory from training history (Fig. 2e, Extended Data Fig. 2b).

Considering the consistent spatial location representations, we further explored whether a shared neural manifold could generalize across subtasks. We found that the subtask spaces were not completely orthogonal. Specifically, the first/smallest principal angles among the 4 go-stimulus-related spaces (AT_Go_in, DT_Go, SR_Go, NR_Go) were significantly smaller than the shuffle levels obtained through random projections o these subtask spaces (Mann-Whitney rank sum test, all p-values << 0.001, Extended Data Fig. 4b). Training a decoder with data from these 4 subtasks allowed it to not only differentiate spatial locations but also generalize to the remaining untrained subtasks (Fig. 2f). Decoding accuracies exceeded shuffle levels (permutation test with Bonferroni correction), while a decoder trained on neuronal data from baseline periods did not exhibit generalization abilities.

What constituted the neuronal basis of this multitasking decoder? By shuffling neuronal activity and assessing contributions to the decoder (Extended Data Fig. 4c), we found that generalization performance across subtasks decreased by approximately 10% when shuffling data from the first 1500 neurons (Fig. 2g). Most of these 1500 neurons were multitasking neurons with mixed selectivity, responding to both task rules and spatial locations. Only 23 location-specific neurons were found among these 1500 (Extended Data Fig. 4c). The high-contribution neurons either maintained their spatial preferences but exhibit different scales in tuning curves (Extended Data Figs. 4d top and 4e top) across subtasks, or exhibited shifted tuning profiles (Extended Data Figs. 4d below and 4e below). These results suggest that a shared multitask neural manifold is widely distributed within the LPFC population, characterized by neurons with mixed selectivity.

Since the LPFC plays an important role in saccadic planning (*2*), we considered whether the shared neural manifold might be influenced by similar eye movement patterns. To address this, both monkeys performed an intercept reach (IR) task (Extended Data Fig. 5a), which had a distinct visual stimulus distribution (Extended Data Fig. 5b) compared to other tasks (e.g. the NR task, Extended Data Fig. 5c), resulting in different touch position patterns (Extended Data Figs. 5d-5f). As the go stimulus was in constant motion, the IR task required the monkeys to track target locations. Thus, the IR task could introduce different eye movement patterns. We then tested whether neural responses from the IR task could be classified within the shared neural manifold (Fig. 2f) across subtasks. The generalization test revealed significant discrimination with ∼40% decoding accuracy (p < 0.001, permutation test), similar to the results from the NR_R1 subtask (Extended Data Fig. 5g). These results confirm that the shared representation of spatial locations across tasks is unlikely to be solely explained by eye movement patterns.

### Meta-task-based representational separations

Having established representational sharing, we next aimed to study representational separation. We sought to ask two questions: 1) Do the same group of neurons separately encode multiple subtasks? 2) If so, are these subtasks equally separated, or grouped into distinct clusters? To evaluate this, we first tested whether all subtasks could be reliably classified. Since neural responses in the SR task were modulated by reward memory (Extended Data Fig. 2b), we subdivided the SR_Go subtask into two based on reward feedback: SR_R (rewarded) and SR_N (non-rewarded). We found that LPFC neurons could indeed differentiate subtasks, as SVM decoders achieved robust classification accuracy for all subtasks (Extended Data Fig. 6a).

To characterize the organization of distinct subtasks, we analyzed dissimilarities between subtask spaces. The geometric dissimilarities included both the orthogonality between pairs of subtask spaces and the distance between their respective offsets. We calculated an orthogonality score for each pair of subtask spaces (Methods), where a score close to 1 indicates high separability, and a score near 0 suggests parallel representations. Our analysis revealed varying levels of orthogonality among the subtask spaces (Fig. 3a, summarized in Extended Data Table 3), with most being separable. The distances between subtask spaces were computed based on the correlation between their offsets in the high-dimensional neural state space. Similar to the orthogonality score, the distances between subtask spaces showed varying patterns of variation (Fig. 3b, summarized in Extended Data Table 4). Thus, although the neuronal representations of different subtasks were separable, the degree of separation varied.

**Fig. 3.**
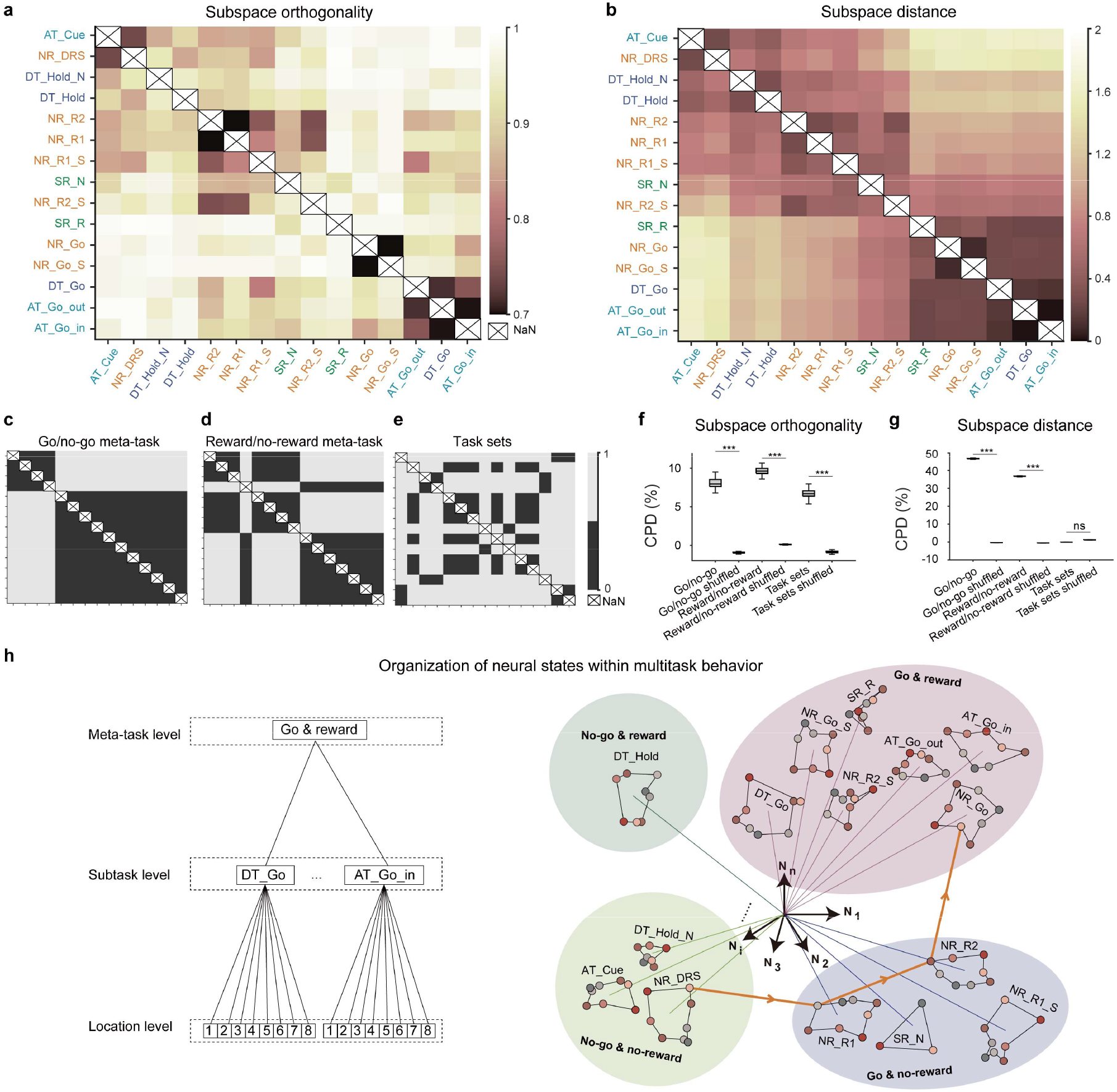
Representational separation of multiple tasks. (**a**) Dissimilarities of space orthogonality. The dissimilarity between each pair of subtask spaces was calculated using the 1 − *VAF ratio* . Font colors on the axes indicate task sets. (**b**) Dissimilarities of space distance. The dissimilarity between each pair of subtasks was measured by Pearson’s correlation distance between the corresponding offset values. (**c-e**) Three templates of representation dissimilarity matrices were constructed, considering go vs. no-go meta-tasks, reward vs. no-reward meta-tasks, and 4 task sets, respectively. The axes across all templates are consistent with **(a)**. (**f-g**) The results from multiple linear regression of the templates onto dissimilarity matrices of space orthogonality (**f**) and distance (**g**). For orthogonality (**f**), CPDs of go/no-go meta-task, reward/no-reward meta-task, and task set were: 53.6 ± 0.1%, 40.5 ± 0.1%, and 1.0 ± 0.0%, respectively (Permutation test, p << 0.001 for meta-task features and p > 0.05 for task-set feature). For distance (**g**), CPDs were: 46.3 ± 0.2%, 36.5 ± 0.2%, and -0.2 ± 0.0%, respectively (Permutation test, p << 0.001 for meta-task features and p > 0.05 for task-set feature). Values are presented as mean ± std. Error bars represent std across bootstraps (n = 100), ***p < 0.001. (**h**) A graphical summary of the hierarchical organization of ∼120 neural states. Left: The hierarchical tree categorized each neural state into three levels: location level, subtask level, and meta-task level. Right: The geometric representation of multiple tasks is organized by this hierarchical tree structure. Hierarchical organization of neural states within multi-task behavior.

We then sought to identify factors that modulate the dissimilarities between subtask spaces. We approached this from two perspectives. The first perspective is the meta-task view, which can group different subtasks into broader meta-tasks. In our task regimes, all the subtasks could be categorized by motor planning (e.g., “plan to go/no-go”) and reward expectation (e.g., “expect reward/no-reward”). According to the meta-task hypothesis, the orthogonality and distance between subtask spaces from different meta-tasks should be larger than those within the same meta-task (Extended Data Figs. 6b-6c). The second perspective is the task-set view, where subtasks are categorized based on their task identities(*13*). Subtasks with the same task identity form a task set, resulting in 4 distinct task sets in our multitask paradigm. If the LPFC utilized the task-set view, the neural representations of subtasks within a task set would be expected to exhibit greater similarity (Extended Data Figs. 6d-6e).

Based on the above hypotheses, we constructed different templates of representation dissimilarity matrices (Figs. 3c-3e). We then applied multivariable linear regressions to quantify the validity of these templates via coefficients of partial determination (CPD) (*34*). The CPD results revealed that the varying levels of subtask space orthogonality could be explained by both meta-tasks (Fig. 3f; go/no-go meta-task, CPD: 8.1 ± 0.6%, permutation test, p << 0.001; reward/no-reward meta-task, CPD: 9.6 ± 0.5%, permutation test, p << 0.001) and task sets (CPD: 6.7 ± 0.6%, permutation test, p << 0.001). However, only meta-task features could explain different levels of distances between subtask spaces (Fig. 3g, CPDs of go/no-go meta-task, reward/no-reward meta-task and task set were: 46.3 ± 0.2%, 36.5 ± 0.2%, and -0.2 ± 0.0%, respectively; Permutation test, p << 0.001 for meta-task features and p > 0.05 for task-set feature). Therefore, the neural representations of subtasks were primarily organized by meta-tasks rather than task sets. These results were consistent across individual monkeys (Extended Data Figs. 7a–b) and individual sessions (Extended Data Figs. 7c– f).

### Hierarchical neural geometry with high generalization abilities

We found that the neural geometry underlying 4 tasks could be conceptualized as a three-level hierarchy comprising ∼120 neural states (Fig. 4a). Each state represented the population-level neural response to a specific spatial location within a subtask. The spatial locations forming a shared ring structure were represented at the foundational level. At the intermediate level, different neural states were grouped by the relevant subtask. At the highest level, subtasks were categorized according to two meta-task features, i.e., motor planning and reward prediction. Consequently, all 4 tasks were represented as distinct subtask sequences in this hierarchical tree structure. To validate this hierarchical organization, we conducted generalization tests across different levels and further explored how this hierarchy could explain monkeys’ behavior during error trials.

**Fig. 4.**
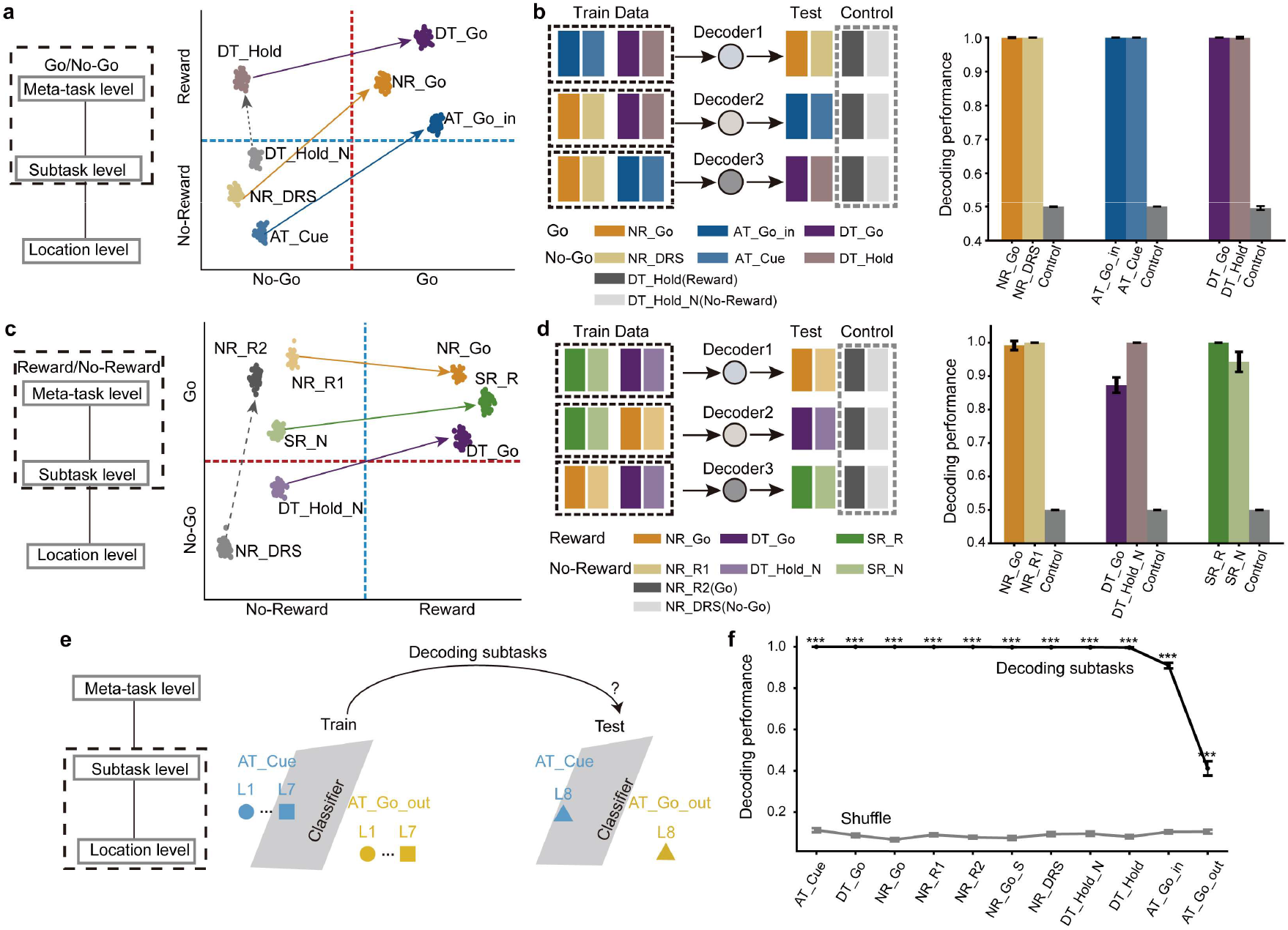
Generalization tests on hierarchical neural representation. (**a**) An illustration of cross-task generalization tests classifying go/no-go meta-task. Left: Generalization tests between meta-task and subtask levels. Right: Illustration of distinguishing multiple subtasks using a shared go/no-go classifier. Each colored dot represents a particular trial from different subtasks in the neural state space. The dashed line represents the classifier, which generalizes across tasks indicated by the arrows for go/no-go discrimination but fails to distinguish reward-related meta-tasks. (**b**) Leave-one-task-out generalization tests for go/no-go discrimination. Left: Testing procedure (Methods). Right: Cross-task generalization results (mean ± std across 100 bootstraps). All performances are above chance level (0.5), except for the control groups. (**c-d**) Similar to (**a-b**), but for reward/no-reward cross-task generalization. (**e**) Leave-one-location-out (LOLO) generalization tests for 11 subtasks. Left: Three-level hierarchical tree showing generalization tests at at the subtask and location levels. Right: Testing procedure (Methods). (**f**) Results of LOLO generalization tests. Values are mean ± std across 100 bootstraps. *p < 0.00455, **p < 0.00091, ***p < 0.00009, permutation test with Bonferroni correction.

If the neural geometry of multiple tasks is hierarchically organized, the neural code at the highest level should remain more abstract and unaffected by changes at the intermediate level. In this case, the neural codes of meta-tasks extracted from subtasks in one task should generalize to subtasks from other tasks. To test this hypothesis, we conducted leave-one-task-out generalization tests for both the go/no-go and reward/no-reward meta-tasks (Figs. 4a-4d). For the go/no-go meta-task, we separately selected a pair of subtasks from each task, excluding the sequential reach task, which contained only the go meta-task (Fig. 4a). If the generalization existed, a classifier trained on data from two tasks should generalize to the remaining task.

Since all subtasks were mixed with both motor-planning and reward-prediction, we additionally used two subtasks assigned to the no-go meta-task category but differing in predicted reward outcomes as a control test set (Fig. 4b left). Neural responses during 330–1000 ms following the task-event onset in correct trials were used to create training and test sets. We trained decoders to discriminate go/no-go meta-task using data from two tasks and tested them on the remaining task (Fig. 4b left). The decoding performance was near perfect across all tests (Fig. 4b right). For the control test, decoders that distinguished reward trials from non-reward trials showed near-chance performance (Fig. 4b right). Similarly, leave-one-task-out generalization tests for the reward/no-reward meta-task (Figs. 4c-4d) demonstrated performance well above chance level, while the decoders failed to distinguish the go/no-go meta-task (Fig. 4d right). These results strongly support a hierarchical relationship between the neural codes of meta-tasks and subtasks.

If the neural geometry was hierarchically structured, the neural code at the intermediate level should remain unaffected by some changes occurring at the lowest level. Thus, we hypothesized that neural codes for subtasks could generalize across different spatial locations (Fig. 4e). We performed leave-one-location-out generalization tests for 11 subtasks (Fig. 4e; three subtasks were excluded due to limited trial number on certain spatial locations) and found that generalization performance across most subtasks was significantly higher than the shuffle levels (Fig. 4f, p < 0.001 for 10 subtasks except for AT_Go_out, which was confused with AT_Go_in).

Could this hierarchical structure also explain the monkeys’ behavior? Given the tree-like organization of the neural geometry, we predicted that neuronal representations at lower levels would be correlated with those at higher levels. Consequently, errors occurring at the subtask level should correlated with errors at the meta-task level across trials (Figs. 5a and 5c). For example, in the DT task, neural responses from false-alarm error trials were decoded as DT_Go rather than DT_Hold at the subtask level, and as “go” rather than “no-go” at the meta-task level (Fig. 5b, averaged neural trajectories of both correct and error trials; Extended Data Fig. 8a, decoding accuracies for both correct and error trials).

**Fig. 5.**
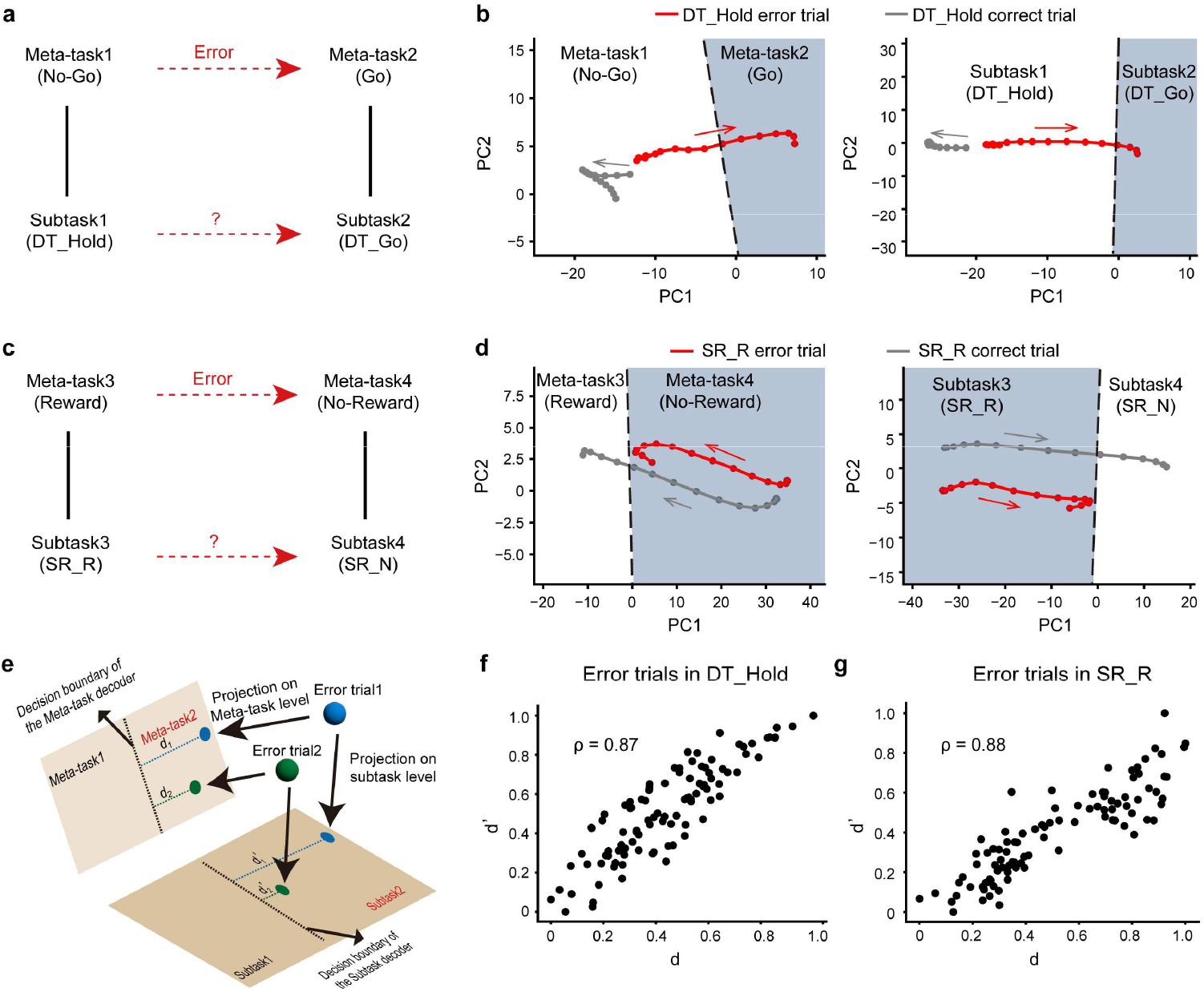
The hierarchical neural representation helps to explain the monkeys’ behavior. (**a**) Errors occurring at the meta-task level (go/no-go) were often observed to correspond with subtask-level errors (e.g., DT_Go/DT_Hold), suggesting a cross-hierarchical relationship. (**b**) The neuronal responses were misclassified at both subtask and meta-task in the DT_Hold error trials. Left: Neural trajectories for correct and FA trials at the meta-task level. The red line represents the average trajectory of false-alarm trials in DT_Hold, while the gray line represents the correct DT_Hold trials. PC1 and PC2 were derived from the PCA analysis on the neural data from NR_DRS, NR_Go, AT_Cue, AT_Go_in trials, with the black dashed line representing the go/no-go decision boundary (Methods). The trajectory of error trials in DT_Hold deviates into the go region, which contrasts with the correct trials. Right: Similar neural trajectories at the subtask level, where error DT_Hold trajectories deviate into the DT_Go region, distinct from correct DT_Hold trials. (**c**) Behavioral errors at reward/no-reward meta-task and SR_R/SR_N subtask levels. Similar to (a), meta-task errors corresponded to subtask errors. (**d**) Misclassified neuronal responses in SR_R error trials. Left: Neural trajectories for correct and error trials at the meta-task level. The red line represents the average trajectory of error SR_R trials, while the gray trajectory represents the correct SR_R trials. PC1 and PC2 were derived from DT_Go, DT_Hold_N, NR_R1 and NR_Go trials, with the black dashed line representing reward/no-reward decision boundary (Methods). The trajectory of error trials in SR_R remains within the no-reward region, contrasting with the correct trials in SR_R. Right: Similar neural trajectories at the subtask level, where error SR_R trajectories remain within the SR_N region, distinct from correct SR_R trials. (**e**) Graphical illustration of the trial-by-trial correlation between the meta-task and subtask levels. (**f**) Trial-by-trial correlation between neural distances to the go/no-go decoder (*d*) and the DT_Go/DT_Hold decoder (*d*′). Scatter plot of distance pairs for error trials, with each dot representing an individual error trial. Pearson’s correlation coefficient is 0.87. (**g**) Trial-by-trial correlation between neural distances to the reward/no-reward meta-task decoder and the SR_R/SR_N subtask decoder, with Pearson’s correlation coefficient of 0.88.

We further computed the distances of error data to decision boundaries of both subtask decoders and meta-task decoders based on correct trials (Fig. 5e). These two sets of distances showed a strong correlation (Fig. 5f, Pearson’s r = 0.87; paired t-test, p < 0.001). To confirm this relationship was not spurious, we performed a control analysis by disrupting the hierarchical relationships through trial index shuffling. This manipulation abolished the correlation between subtask and meta-task levels (Extended Data Fig. 8b). This cross-level correlation was not restricted to any specific task. In the SR task, monkeys made errors in reward predictions by touching incorrect fourth-rank locations. These errors were associated with misclassifications in both subtask and meta-task decoders (Fig. 5d, averaged neural trajectories of both correct and error trials; Extended Data Fig. 8c, decoding accuracies of both correct and error trials). As with the DT task, the distances of error data to decision boundaries for both subtask and meta-task decoders were highly correlated (Fig. 5g; Pearson’s r = 0.90; paired t-test, p < 0.001). As a control, no correlation was observed when disrupting the relationships between the subtask and meta-task levels (Extended Data Fig. 8d). These findings suggest that monkeys’ error responses at certain subtasks can be explained by the neural codes of meta-tasks.

### Anatomical organization of neural code for multitask in LPFC

The utilization of two-photon imaging allowed us to explore the anatomical arrangement of neural codes across different levels of the hierarchy. We first assessed whether the neurons with similar location preferences were anatomically closer. For the neurons in each FOV, we calculated clustering indices at the location level based on the regression coefficients from the same subtask, and compared these indices to shuffled distributions obtained by randomly permuting the positions of all neurons within the same FOV (Fig. 6a, Methods). To better explore the anatomical organization of location codes within each FOV, we selected the subtask with the largest location variability (Figs. 6a and 6d, NR_Go for the example FOV). Our analysis revealed a mild but significant anatomical clustering of location preference at the scale within 150 μm for the subtask exhibiting the strongest spatial codes (Fig. 6g, an example FOV, permutation tests, p < 0.0001 on “0-50 μm” group and “100-150 μm” group; Fig. 6h, all FOVs “0-50 μm” group; Extended Data Figs. 9a-9f, results for individual monkeys).

**Fig. 6.**
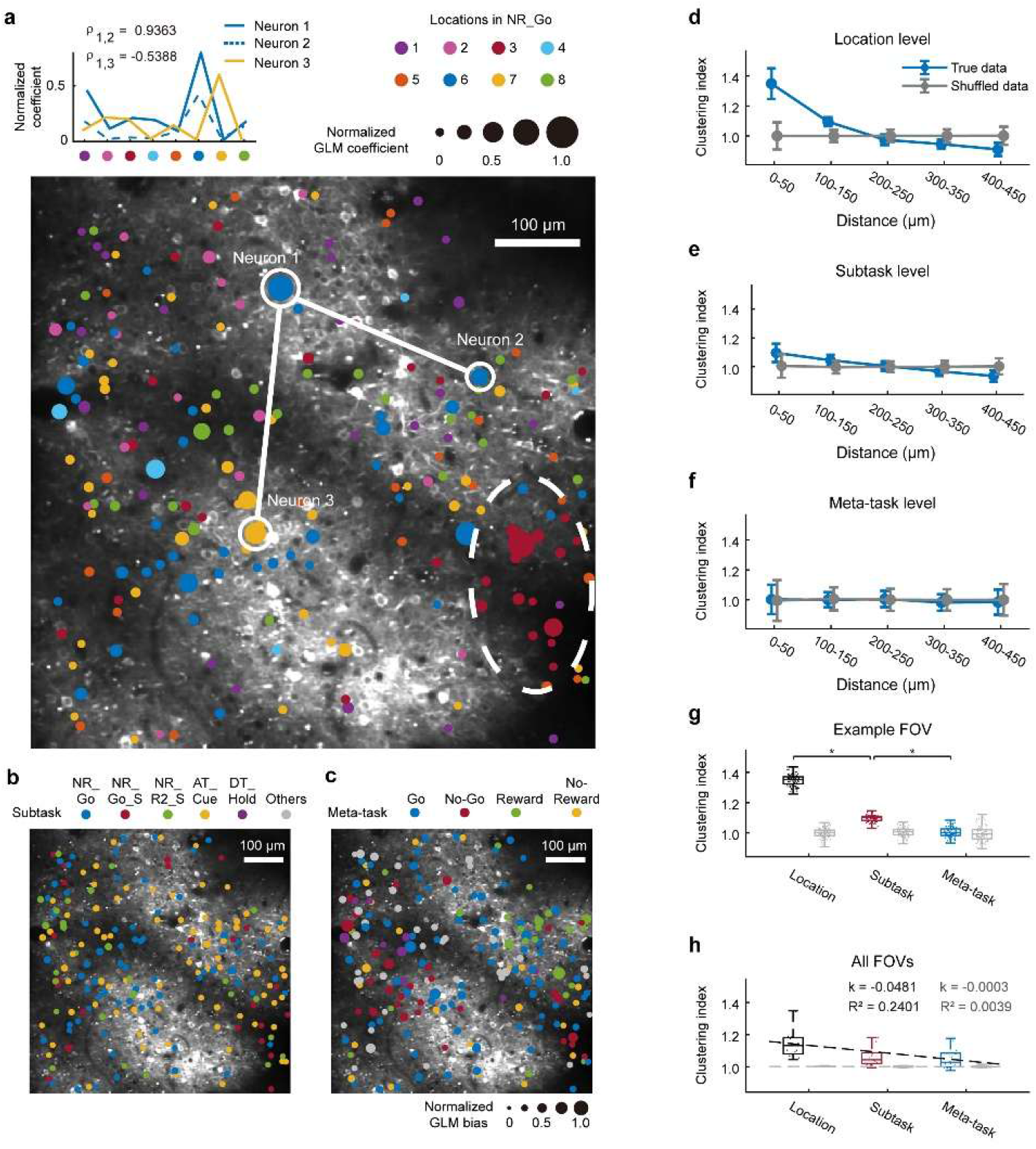
The anatomical organization of representations at different hierarchies. **(a)** Location tuning of LPFC neurons at FOV 05 from monkey 1. Filled circles represent ROIs, where colors indicate preferred locations based on GLM coefficients, and sizes reflect normalized GLM coefficient values (Methods). Neurons preferring location 3 were predominately clustered in the lower-right subregion highlighted by a white dashed ellipse. GLM coefficients for example neurons are shown, with correlation coefficients (*ρ*) between neurons calculated from these values. Cluster indices were determined from correlations between neuronal pairs within specific distance ranges (Methods). (**b-c**) Similar analysis as in (**a**), but with normalized GLM bias values for subtasks (**b**) and meta-tasks (**c**) respectively (Methods). (**d-f**) Anatomical clustering analysis results for location (**d**), subtask (**e**), and meta-task (**f**) levels. Neurons with small distances (0-50 μm) exhibited larger similarity at the location level indicated by blue curves, which was significantly higher than the shuffled results indicated by gray curves (permutation test, p < 0.0001). Long-range pairs (300-450 μm) had lower clustering than shuffled data (permutation test, p < 0.0001). Weaker but significant clustering effects were observed at subtask level within 50 μm (permutation test, p < 0.0001). No anatomical clustering was observed at the meta-task level. X-axis: distances between neuronal pairs; Y-axis: clustering index. Error bars indicate 5 × std. (**g**) Clustering indices of 0-50 μm neuronal pairs across levels in an example FOV. Each dot representing a bootstrap resampling (Methods) is color-coded by the hierarchy. Gray boxes and dots indicate shuffle levels. The clustering indices significantly decreased with increasing hierarchical levels (all p-values < 0.0001, permutation tests). (**h**) Clustering indices at 0-50 μm scale across all FOVs. Similar analysis and format as (**g**). The black dashed line indicates linear fit of clustering indices across levels (*k* = 0.0481, *R* = 0.2401). The gray dashed line indicates shuffled data (*k* = 0.0003, *R* = 0.0039).

We then extended above analysis to investigate further anatomical organizations of the neural codes at subtask (Figs. 6b and 6e) and meta-task levels based on the regression offsets (Figs. 6c and 6f).The clustering analysis still indicated weak anatomical organization at subtask level at the scale of < 150 μm (Fig. 6e, FOV 05, permutation tests, p < 0.0001 within 50 μm). But there was no clustering effect at meta-task level (Fig. 6f). Therefore, our results indicate that anatomical organization of neural codes in the LPFC follows a decreasing trend across increasing hierarchical level (Fig. 6g, example FOV, permutation tests, p < 0.0001 between location and subtask levels, p < 0.0001 between subtask and meta-task levels; Fig. 6h, all FOVs; Extended Data Figs. 9g-9h, individual monkeys).

## Discussion

By employing two-photon calcium imaging on the same LPFC neurons while monkeys perform multiple tasks, we uncovered the representational geometry of these tasks within the LPFC neural state space. Both shared and separated representations were identified. The neural codes for spatial locations within each subtask exhibited similarity within their respective low-dimensional spaces, revealing a generalizable neural manifold. Although the neural representations of subtasks were distinct, the varying levels of dissimilarity between subtask spaces could primarily be attributed to meta-task characteristics. Therefore, the neural states of task variables were organized hierarchically: the neural representations of spatial locations were embedded within subtask spaces, which were further grouped by meta-task categories. This hierarchical structure supported generalization across different tasks and locations, and could further help explain monkeys’ behavior on error trials. Furthermore, the anatomical clustering of these representations showed an inverse relationship with hierarchical level, decreasing as the level of abstraction increased.

Hierarchical organization is essential to a range of cognitive functions (*35-37*), including cognitive control (*38*), working memory (*39*), language (*40*), and concept learning (*41*). Our findings suggest this organization can also support both representational sharing and representational separation across different tasks. In terms of representational sharing, spatial locations were encoded similarly across subtasks within low-dimensional subspaces, forming a ring-like structure. We further found that a shared neural manifold for encoding spatial locations could generalize across various tasks. This representational sharing extended beyond spatial locations, as our generalization tests showed that hierarchical structure could be applied across locations and tasks. These results imply that shared representations facilitate efficient generalization, potentially reducing learning costs for novel tasks by leveraging preexisting neural codes. Further research could explore the role of this hierarchy in rapid task acquisition and transfer learning. Effective multitasking, however, requires not only shared representations but also clear separation between tasks to minimize interference and maintain behavioral performance. We found that, although many neurons exhibited mixed selectivity for spatial locations and task rules, subtask spaces retained distinct separations, characterized by orthogonal bases and separate offsets. The separation of subtask spaces ensures distinct neural encoding across different tasks, therefore minimizing interference and enabling precise task execution.

The tree-like structure of this representation allows for separation at lower levels while promoting shared representations at higher levels. This hierarchical geometry integrates both representational sharing and separation, supporting cognitive flexibility while maintaining task stability.

Both abstract task structures and particular sensory objects emerge in the brain. Factorized representations not only support representational separation (*32, 42*), but also enable preexisting structural knowledge to seamlessly guide learning (*43-45*). Previous research has identified two neuron types involved in multiple tasks: task-specific neurons (*14, 15*) and sensory-rule mixed neurons (*46*). These findings complicated our understanding of how individual neurons encode multiple tasks.

In our study, we demonstrated that both mixed neurons and task-specific neurons can be effectively described by the same regression models. The regression coefficients encode a mixture of sensory-related and task-related information, while the bias terms reflect task-specific information. By concatenating regression coefficients and bias terms across all neurons, we vectorized task-related variables within a high-dimensional neural state space. The offset of each subtask space was characterized by the regression biases. In this state space, the spatial locations that occurred in each subtask were embedded in the corresponding low-dimensional space. The representations of spatial locations exhibited a shared ring-like structure across different subtask spaces, suggesting a task-free neural code for sensory input. Conversely, the orthonormal basis and offset of each subtask space remained invariant to spatial location. Therefore, a factorization of sensory-related and task-rule-related neural codes emerged at the collective level. This suggests a unified view of multitask representation at both single-neuron and population levels, shedding light on how neurons integrate sensory information with abstract task rules to coordinate flexible, goal-directed behavior.

In the present study, we designed a multitask behavior paradigm involving tasks related to go/no-go decision making (*47, 48*), selective attention (*49, 50*), reward-related memory (*51, 52*), and dual task (*53*). To streamline training, we simplified these tasks. Although the eye movements and hand movements were not tightly controlled, the behavioral results and neural activities of both monkeys aligned with previous findings, indicating successful implementation of all four cognitive processes. The LPFC is recognized for its central role in a wide range of cognitive functions (*17, 54*), and thus serves as a hub for multiple tasks(*55-57*). However, few studies have investigated how these cognitive functions are encoded by the same group of neurons. Our results illustrate a neural geometry comprising both shared and separated modules, providing insight into the operational mechanisms of this hub. While the selected tasks were designed to reflect specific cognitive processes, we acknowledge that they may not fully capture the relationships among different cognitive functions. Future research should expand on this framework by exploring a broader range of cognitive processes.

Over 70 years ago, Karl Lashley proposed that complex sequential behavior would be controlled by a hierarchically organized program (*58*). In line with Lashley’s insight, our study shows that multitask behavior, consisting of subtask sequences, emerges from a hierarchical tree structure, offering a fundamental neuronal mechanism for both cognitive flexibility and stability.

## Resource availability

## Data and code availability

All data and code used in this study are available from the corresponding author upon reasonable request.

## Acknowledgments

We would like to thank Liping Wang, Tianming Yang, and Bin Min for their valuable comments on the manuscript and analyses.

## Funding

Shanghai Rising-Star Program 22QA1412100 (Y.X.)

Lingang Laboratory startup fund (Y.X., Z.C.)

National Key R&D Program of China 2021ZD0203601 (C.L.)

National Key R&D Program of China 2019YFA0709504 (C.L.)

Shanghai Municipal Science and Technology Major Project 2021SHZDZX (C.L.)

National Natural Science Foundation of China 32221003

National Natural Science Foundation of China 32161133024

Shanghai Pilot Program JCYJ-SHFY 2022-010 (C.L.)

Lingang Laboratory LG-GG-202402-06 (C.L.)

## Author contributions

Conceptualization: C.L., Y.X., Z.C.

Methodology: Z.F., Y.X., Q.S., S.L., J.J., D.L., Z.H., Y.C., Y.S., F.W.

Investigation: Z.C., C.L., Y.X.

Visualization: Y.X.

Funding acquisition: Z.C., C.L., Y.X.

Project administration: Y.X.

Supervision: Z.C., C.L., Y.X.

Writing – original draft: Y.X.

Writing – review & editing: Q.S., S.L., D.L., J.J., Z.C., C.L., Y.X.

## Competing interests

The authors declare no competing interests.

## SI

### Experimental animal model and surgical procedures

Two male cynomolgus macaques (*Macaca Fascicularis*) were used in the study. Two monkeys (M1 and M2, 4.4 kg and 4.3 kg, 14 years old) were used for two-photon calcium imaging. All experimental protocols of the two-photon imaging study followed the Guide of the Institutional Animal Care and Use Committee (IACUC) of Institute of Neuroscience, Chinese Academy of Science (CEBSIT-2021068R01).

Two surgeries were performed on each animal under general anesthesia. In the first surgery, head posts were implanted onto each animal’s skull. Two head posts were implanted on the back of the head and one on the forehead. A Y-shaped steel frame was fixed to these head posts for head stabilization during training and recording.

After reaching the performance criteria, the second surgery was performed. In the second surgery, a 22 mm craniotomy was made on the skull over the region of lateral prefrontal cortex located along the principal sulcus (PS) reaching the arcuate sulcus (AS), roughly corresponding to Walker’s areas 46 and 8(*59*), and the durotomy (diameter 16 mm) to expose the cortex. Then, the virus injections were performed. The mixture of around 250 nL AAV2/9.pAAV.TRE.GCaMP6f (titer: 6e12 GC/ml, Gene Editing Core Facility, Institute of Neuroscience) and 250 nL AAV2/9.pAAV.hSyn1.tTA (titer: 2e12 GC/ml, Gene Editing Core Facility, Institute of Neuroscience) was injected into the tissue at a depth of around 500 μm(*60*). The injection sites were located alongside the PS, roughly corresponding to Walker’s area 46 (Extended Data Fig. 1). After virus injection, an imaging window was implemented. A glass coverslip (18 mm in diameter and 0.2 mm in thickness) glued with a titanium ring (15 mm outer diameter and 13 mm inner diameter) constituted a window unit. The window unit was gently pushed onto the cortical surface. The coverslip was inserted under the dura and glued with medical adhesive. The titanium ring was glued to the skull with dental acrylic. The entire imaging window was then covered with a steel shell to protect the coverslip. For further details, please refer to Li et al. 2017(*61*).

### Behavioral task

Two monkeys were trained to complete four tasks where all visual stimuli and motor outputs were associated with the same eight spatial locations that arranged in a square formation, respectively. Each task comprised multiple serially organized subtasks. And a specific subtask is related to a particular task event. The monkeys were seated in a primate chair facing a 21-inch LCD touch screen at a distance of one arm length around 30 cm during experiments, the front of the chair was 15 ∼ 22 cm away from the screen. A computer-controlled solenoid determined the reward size. We used Matlab psychtoolbox to control behavior and collect behavioral data.

#### The behavioral tasks were detailed as follows

##### Discrimination task

During each trial, the monkeys were tasked with discriminating between two types of visual stimuli displayed on the touch screen. When the stimulus was a mole which referred to as “go trials”, the monkeys need to reach its location. The eccentric range of a valid hit was 2.5 cm around each location in the go trials. Conversely, when the stimulus showed an owl or a snake which referred to as “no-go trials”, the monkeys were instructed not to touch its location. In the go trials, correctly touching the designated locations within 330-4000 ms were classified as hit trials, while the rest trials were categorized as “miss trials”. The time limit for each trial was uniformly distributed random values within a range, which varied among different sessions (Extended Data Table 1). On the other hand, in the no-go trials, touching the owl or snake was considered a “false alarm trial”, while refraining from touching led to “correct rejection trials”.

Various images corresponded to different types of feedback in the experiment. In instances where monkeys made accurate choices, successful selections involving either the mole or owl images triggered the dispensing of a single drop of juice, serving as positive reinforcement. Conversely, no reward was given for selections associated with the snake image. In the error trials, missing responses led to no reward, while false alarms prompted the display of full-screen images for approximately 1 second without any reward as negative feedback. Specifically, during a trial featuring an owl or snake, a full-screen owl or snake image was presented, respectively. Collectively, all trials in discrimination task could be categorized into three groups: go with reward, hold with reward, and hold without reward (a 1:1:1 ratio).

##### Attention task

During each trial, the monkeys were tasked with touching the target’s location within a time window of 300-670 ms (Extended Data Table 1). The target was depicted as a mole image followed by a hint cue in the form of a red dot briefly shown for 200 ms at one of eight candidate locations. The task was divided into two types according to whether the cue’s position was consistent with the target position. In 75% of all trials, the hint cues aligned with the target locations, referred to as “cue-in trials”, while in the other 25% of all trials, they did not align, referred to as “cue-out trials”. A single drop of juice was dispensed upon successful completion of each correct trial. Notably, the monkeys’ eye movements were unrestricted, and the attention modulation in this task was considered as overt attention(*62*).

##### Sequential reach task

During each trial, monkeys were tasked with sequentially reaching four locations to form a length-4 reach sequence. The target locations were indicated by mole images, and the monkeys had to respond within 330-670 ms on each ordinal rank. Upon successfully completing a trial, two drops of juice were delivered. If an error was made during a trial, the trial ceased immediately. In 80% of the trials, the reach sequence followed the pattern of 1-3-5-7, while in the remaining trials known as “catch trials”, the target on the final ordinal rank was randomly assigned to one of seven locations other than location 7.

##### Nested reach task

The nested reach task comprised an outer task of a delay reach and an inner task of two consecutive reaches. The distribution of inner task, outer task, and nested task was 10%, 20%, and 70%, respectively.

In the outer task trial, a sample cue appeared at one of eight locations for 1000 ms. After a 1000 ms delay period, a go signal prompted the monkeys to reach the sample cue location within a specified time frame. The sample cues, indicating delayed responses, were labeled as delayreach stimulus (DRS) cues. Successful completion of an outer task trial within 4000 ms led to the delivery of two drops of juice.

In the inner task trial, monkeys were tasked with reaching two locations consecutively, guided by mole images, within 500-4000 ms (Extended Data Table 1) for each response. A correct response in the inner task trial resulted in the delivery of one drop of juice.

In the nested task trial, monkeys had to complete the inner task correctly after the DRS cue presentation. Successful completion of a nested task trial within 4000 ms led to the delivery of two drops of juice. Any error during a trial led to an immediate halt of the trial.

##### The abbreviations of task names

The abbreviations of task names and subtask names in all four tasks are detailed below, and the subtask name is the same as the corresponding task event:

DT: Discrimination task.

DT_Go: Go stimulus in DT task.

DT_Hold: No-go and predicting-reward stimulus in DT task.

DT_Hold_N: No-go and predicting-no-reward stimulus in DT task.

AT: Attention task.

AT_Cue: Attention cue in AT task.

AT_Go_in: Attention stimulus whose location was consistent with attention cue.

AT_Go_out: Attention stimulus whose location was inconsistent with attention cue.

SR: Sequential reach task.

SR_Go: Go stimulus in SR task, including both SR_R and SR_N.

SR_R: Predicting-reward stimulus in SR task (i.e. the 4th rank in both normal and catch trials).

SR_N: Predicting-no-reward stimulus in SR task (i.e. the first 3 ranks).

NR: Nested reach task.

NR_R1_S: First reach in standalone inner task.

NR_R2_S: Second reach in standalone inner task.

NR_Go_S: Go signal in standalone outer task.

NR_DRS: Delay reach stimulus in nested outer task.

NR_R1: First reach in nested inner task.

NR_R2: Second reach in nested inner task.

NR_Go: Go signal in nested outer task.

### Behavioral analysis

All tasks shared the same type of trials that required monkeys to reach to the correct locations. To measure monkeys’ performances under these shared conditions, we calculated the proportion of correct trials on each location, and grouped monkeys’ performances across all imaging sessions.

Apart from the shared conditions, we also targeted different cognitive processes underlying each task, and measured whether and how these cognitive processes influenced monkeys’ behavior.

#### Discrimination task

To assess the monkeys’ ability to distinguish between two types of visual stimuli (go vs. no-go cues) and respond accurately, we compared the proportions of responses in the go trials (hit rate) and the no-go trials (false-alarm rate) with paired samples t-tests.

#### Attention task

To investigate if attention cues could effectively direct the monkeys’ attention, we compared the monkeys’ reaction times (RTs) between the cue-in group and the cue-out group. The RT was defined as the time of screen touch minus the time of the go signal. RTs for the same locations in each group were organized alongside imaging sessions, and a paired samples t-test was conducted to evaluate whether attention cues significantly modulated the monkeys’ attention.

#### Sequential reach task

If the monkeys had mastered this task, they would be able to anticipate the locations of the go stimulus for each ordinal rank. Thus, there would be a discrepancy between the catch trials—where the rank-4 locations were not on location 7—and the normal trials, which went against the monkeys’ predictions and resulted in longer reaction times. As a control, the locations on the first three ranks of catch trials would show no difference compared with those of normal trials. Therefore, we anticipated that only rank 4 exhibited longer RTs when catch trials were compared with normal trials. RTs for each rank from catch trials or normal trials were organized alongside sessions, and paired samples t-tests were conducted to examine the existence of differences between normal and catch trials.

#### Nested reach task

We calculated correct rates and reaction times of both monkeys for the inner task trials, the inner task trials within nested task, the outer task trials, and the outer task trials within nested task. To further evaluate interference from the inner task on the outer task, we calculated the proportion of trials where the monkeys’ choices in the outer task were affected both by the first and the second reach location in the same trials. We grouped the error proportions from all trials across sessions and conducted Wilcoxon signed-rank test to determine whether these error proportions were significantly higher than the chance level. For the outer task, with 8 candidate locations, the chance level was set at 1/8. Monkey 1’s errors during the subtask NR_Go were significantly located on the inner task targets (R1/R2, Fig. 1f left), and meanwhile, monkey 2’s incorrect responses might still cluster around them, which could also reflect a subtle inner task interference. To test this, we compared the distribution of distances between errors and R1/R2 with a chance distribution (Extended Data Figs. 1b-1c).

### Two-photon calcium imaging and data preprocessing

We performed in vivo two-photon calcium imaging using a Thorlabs two-photon microscope BERGAMO II and a Ti: Sapphire laser (Chameleon Vision S, Coherent). A 16 × objective lens (0.8 N.A., Nikon) imaged an area of 512 μm × 512 μm or 800 μm × 800 μm at 30 frames per second. The recording depth ranged from 200 μm to 350 μm below the pia. Each imaging session lasted for 2 ∼ 4 hours.

The image processing pipeline was implemented in Python and JupyterLab(*32, 63*). First, two-photon images were temporally down-sampled and spatially smoothed with a Gaussian filter. Then, the images were motion corrected with non-rigid algorithms. When the motion artifacts were too severe, these imaging data were excluded after examining the spatial shifts and correlations with the reference image.

Source extraction was performed with the CaImAn package(*63*) based on constrained non-negative matrix factorization. A set of scores were calculated for each extracted component. Regions of interest (ROI) were selected by either thresholding these scores or, in ambiguous cases, human inspection. The resulting traces had a frame rate of 15 Hz. The calcium traces were first detrended and then normalized by the noise variance, which was estimated by calculating the variance of the residual signal after subtracting a Savitzky-Golay filtered version. There were 16 recording sessions as well as FOVs in total (12 for M1 and 4 for M2). Around 200 ROIs on average were found in the whole corresponding imaged FOVs (ROI number in each FOV, Extended Data Table 2). These FOVs were distributed alongside the PS, primarily located at posterior and ventral sides (Extended Data Fig. 1g).

### Single neuron analysis

To better compare single neuron activities across spatial locations, we selected subtasks with similar visual input and motor output across all tasks. These include the epoch when the go stimulus was presented (DT_Go), the epoch when the location of the go stimulus matched the attention cue (AT_Go_in), the epoch in the sequential reach task (SR_Go), and the epoch in the nested reach task (NR_Go). All epochs were selected from the 0-2000 ms window after the onset of corresponding subtasks. Additionally, to analyze neuron responses before task onset, we chose the data during 0 ∼ 1000 ms before trial onset as baseline for all four tasks. For simplification, neural data from the baseline period and the task epochs within the same trial were concatenated.

To categorize each individual neuron, we first averaged ∼500 ms (8 frames at 15 Hz sampling rate) of neural responses after the relevant task onset within each trial, and then conducted two-way ANOVA for 8 locations and 4 tasks. Therefore, we obtained variances attributable to the task-related factor (i.e., the location factor, task factor and locations × tasks mixed factor)) for each neuron. Then, we shuffled task and location labels 1000 times to generate a distribution of label-shuffled variance and calculated the probability *p*_*k*_ that variance with true labels was smaller than label-shuffled variance, respectively:

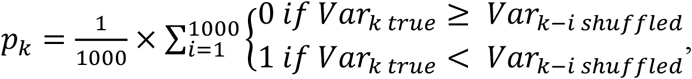

in which *k* is the task-related factor and *i* is the count of shuffling.

We defined neurons with p < 0.05 for location factor and other p-values >= 0.05 as location-specific selective cells, whereas neurons with p < 0.05 for task factor and other p-values >= 0.05 are task-specific selective cells, and neurons with p < 0.05 for both location and task factors or the interaction factor as mixed-selective neurons. The remaining neurons were classified as non-selective cells.

Among mixed-selective neurons, we applied one-way ANOVA to estimate the location selectivity in each task following the same protocol. We considered that these mixed-selective neurons contributed to the locational encoding on tasks in which p-value < 0.05. As a result, mixed-selective neurons were categorized into single-tasking neurons and different levels of multitasking neurons (25 mix neurons contributed to none of four tasks, which resulted from the lower trial number in single task analysis; those neurons were excluded from the proportional statistics).

### Neural state space analysis

In the analysis of the neuronal population, we initially collected all ROIs from 16 FOVs to create a pseudo-population. Within this high-dimensional neural space, a state, as defined in our experimental setup, represented a combination of a specific location *l* and subtask *t*. By gathering the states from the baseline periods of all four tasks, we observed 14 subtasks in our multitask paradigm. Specifically, where in Figs. 2f-2g, 3 subtasks originated from the DT task (DT_Go, DT_Hold, and DT_Hold_N); 3 from the AT task (AT_Cue, AT_Go_in, and AT_Go_out); 1 from the SR task (SR_Go); and 7 from the NR task (NR_DRS, NR_R1, NR_R2, NR_Go, NR_Go_S, NR_R1_S, and NR_R2_S).

### Linear regression

Then we fitted a general linear regression model (GLM) to determine how various task variables affected the average neural response in each subtask. Neuronal preferred locations within a specific subtask can be represented as an 8-dimensional one-hot vector. We thus defined 8 one-hot vectors *β*_*t,l*_ as each subtask variables, where *t* = 1, ⋯, 14 and *l* = 1, ⋯, 8. In our model, for a given subtask *t*, the averaged neural response of the *i*-th neuron in one trial was assumed to be a linear combination of subtask variables, such that:

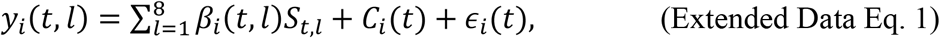

where *β*_*i*_(*t, l*) are unknown regression coefficients and *C*_*i*_*(t*) is the bias term and *ϵ*_*i*_*(t*) is the trial-by-trial noise. To avoid overfitting, we resampled 80% trials and incorporated an L1 regularization term into the GLM framework, with the regularization strength optimizing through a 5-fold cross-validation and minimizing the mean squared error. We applied each model to 100 bootstraps resampled datasets.

Specifically, when subdividing SR_Go into SR_R and SR_N, we conducted two additional GLMs correspondingly. Notably, the SR_N condition utilized a 3-dimensional one-hot vector due to its systematical restriction to only three spatial locations.

### Targeted dimensionality reduction (TDR)

Targeted dimensionality reduction was a methodological framework, which contains grouping neuronal data with shared features and dimensionality reduction(*25, 29, 32*). Following this method, we identified a group of subspaces, each of which captured major variance of the neural states within the same subtask. And these subspaces were termed subtask spaces.

### Geometric view of state space analysis

A state, in our task regime, could be vectorized as an *N*-dimensional vector in the neural state space. This vector could be decomposed into two hyperparametric vectors. The first one was embedded in a low-dimensional subtask-*t* space defined by the orthonormal basis ***Q***_***t***_, resulting in a 2-dimensional projected value ***f***(*t, l*) that related to both the subtask and spatial location. The second one was the *N*-dimensional biased vector ***C***(*t*), solely indicating the center of the subtask-*t* space. Consequently, the subtask spaces ***Q***_***t***_, ***f***(*t, l*) and the biased vectors ***C***(*t*) were utilized to characterize the geometric neural representation of multiple tasks.

### TDR on identifying regression-based subtask spaces

The regression coefficients *β*_*i*_*(t, l*) were used to identify the low-dimensional subspaces which captured major variance of the neural states from same subtask. Specifically, with *N* neurons in total collected from all FOVs, an *N*-dimensional vector ***β***(*t, l*) = [*β*_1_(*t, l*), ⋯, *β*_*N*_*(t, l*)]^*T*^ was defined as the populational representation of the state with subtask *t* and location *l*. To capture the response variance induced by location variation at each subtask in this neural state space, principal component analysis implemented via singular vector decomposition (SVD) method was performed on each group {***β***(*t, l*)} _l=1,⋯,8_ to identify the first two axes that captured the most response variance (see Extended Data Fig. 2c for the cumulative explained variance). Therefore, with PCA analysis, we approximated the neural state space as:

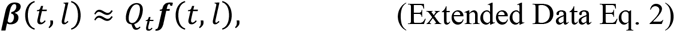

where 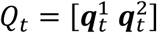 with 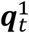 and 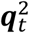 forms an orthonormal basis for subtask-*t* space, ***f***(*t, l*) is the projected value on this subtask-*t* space under the orthonormal basis 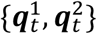 . In other words, we obtained the collective variable ***q***(*t, l*) by projecting the demeaned ***β***(*t, l*) onto subtask-*t* space.

*C*_*i*_*(t*), the bias of the regression model was used to identify the low-dimensional subspaces containing variance at subtask level. Specifically, an *N*-dimensional vector ***C***(*t*) = [*C*_1_(*t*), ⋯, *C*_*N*_*(t)]*^*T*^ ∈ ℝ^14×*N*^ was used to measure subtask information at the neuronal population level. From the geometric view, vector ***C***(*t*) specified the centroid of the subtask *t* space. Therefore, ***C***(*t*) could measure the distance from the centroid of each subtask space to the origin of the neural state space. To capture the variabilities of neural responses induced by subtask variation in this neural state space, principal component analysis implemented via SVD method was performed on all ***C***(*t*) to identify the first two axes that captured the most variance (see Extended Data Fig. 2d for the cumulative explained variance).

### Behavior relevance of subtask space analysis

Discrimination task (Extended Data Fig. 3a): To illustrate the relationship between neuronal representation and the behavioral relevance in DT task, we first determined the DT space by conducting PCA on the matrix ***C***(*DT*) ∈ ℝ^3×*N*^, which was obtained by concatenating the offsets for each subtask (DT_Go, DT_Hold, and DT_base as the newly GLM offsets) in the discrimination task, where *N* denotes the number of neurons with error trials. Then we aligned the behavioral-related neuronal trajectories by averaging across 330-1000 ms following the onset of the go/no-go stimulus of each correct trial per neuron, to obtain 2-dimensional matrices *M*_*DT_Hold*_ ∈ ℝ ^*K×N*^ and *M*_*DT_Go*_ ∈ ℝ ^*K×N*^. As for DT_base, the neuronal activities were averaged across 100-800 ms prior to stimulus onset to obtain the matrix *M*_*DT_Base*_ ∈ ℝ ^*K×N*^ . The three matrices were then projected onto DT space to determine the distribution of each subtask. With the false alarm trials in the discrimination task, we constructed the error neuronal activities *M*_*DT_FA*_ ∈ ℝ ^*K×T×N*^ with the pseudo-population method and then averaged across the trial dimension to obtain the mean error temporal trajectory *M*_*DT_FA_Mean*_ ∈ ℝ ^*T×N*^. To smooth the trajectory, we applied a sliding window of length 660 ms and step size 66 ms along the temporal dimension, subsequently projected onto the DT space as well. As a control, we obtained a mean correct temporal trajectory *M*_*DT_CR_Mean*_ ∈ ℝ ^*T×N*^ by averaging the correct rejection trials following the same protocol. To verify the results based on the method mentioned above, we also used matrices *M*_*DT_Hold*_ and *M*_*DT_Go*_ to train a decoder to discriminate between DT_Go and DT_Hold. Decoder performance was evaluated by contrasting trajectories from false alarm trials with those from correct rejection trials, with both datasets averaged across the 330-1000 ms time window following stimulus onset.

Attention task (Extended Data Figs. 3b-3c): In a comparative analysis of the neural response between attend-in and attend-out trials, we first combined regression coefficients of all neurons from subtasks AT_Go_in and AT_Go_out, which was denoted by a matrix *M*_*AT*_ ∈ ℝ^*100×16×N*^, where *N* denotes neuron number, 16 denotes 8 locations from two subtasks, 100 denotes bootstrapping results mentioned in “Linear regression” section. Then, we identified an averaged 2-D space by averaging 100 bootstrapping results to get a matrix 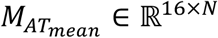 and performed PCA on it. In this way, we illustrated the neural geometry of the neuronal representation with different levels of attention by projecting each geometry of AT_Go_in and AT_Go_out onto this subspace, and measured the sizes of two geometries in the neuronal representation with the shoelace formula. To compare with the shoelace sizes of different population from all FOVs and the single FOV, we conducted the identical operations in each FOV. Then we applied min–max normalization to individual FOV geometries to eliminate inter-FOV baseline variations while preserving relative spatial patterns.

Nested reach task (Extended Data Fig. 3d): To illustrate the interference of dual samples in the nested reach task, we conducted PCA analysis on the matrix ***C***(*NR*) ∈ ℝ^*4×N*^, which was obtained by concatenating the offset for each subtask, to determine the 3-D NR space, where 4 denotes four subtasks (NR_DRS, NR_R1, NR_R2 and NR_Go) and *N* denotes the number of neurons included in error trials. We projected three matrices *M*_*NR_DR*_ ∈ ℝ ^*K×N*^, *M*_*NR_R*_ ∈ ℝ ^*K×N*^ and *M*_*NR_Go*_ ∈ ℝ ^*K×N*^, which constructed using pseudo-population method and then averaged across the 330-1000 ms following stimulus onset, onto the NR space, to determine the distribution of each subtask within the NR space. With the trials in which monkey selected false locations of DRS cue, we constructed the error neuronal activities with the pseudo-population method and then averaged across the trial dimension to obtain the mean error temporal trajectory. We smoothed the trajectories following the same protocol as introduced in the discrimination task, subsequently projected onto the NR space. As a control, we obtained a mean correct trajectory with the correct trials using the same protocol. We also used matrices *M*_*NR_R*_ and *M*_*NR_Go*_ to train a decoder to discriminate the second nested reach stimulus and go signal. The decoder was then tested using the averaged neuronal trajectory data averaged from 330-1000 ms after stimulus onset of the false trials and the correct trials as control.

### Similarity analysis of representational geometries

To analyze the neural representation of 8 spatial locations, we sought to understand the degree of similarity among all the geometric representations within each subtask space. Specifically, we approximated collective variables ***f***(*t, l*) with an affine transformation, such that

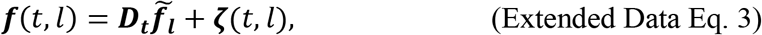

where ***D***_***t***_ is

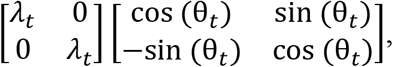

combined with scale transformation *λ*_*t*_ and rotation transformation 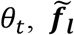 is a spatial location vector independent of subtask *t* and ***ζ***(*t, l*) is the approximation error. In practice, we obtained ***D***_***t***_ and ***f***_***l***_ through minimizing the Frobenius norm of the error matrix 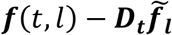 while maximizing the Frobenius norm of matrix 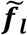,

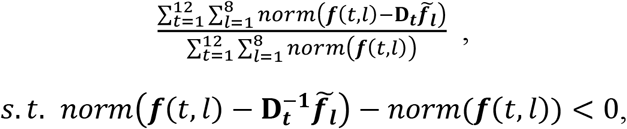

by leaving SR_Go and NR_Go out as testing sets, we selected 12 subtasks to get 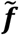, and calculated the similarity score between 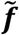 and the ***f***(*t*,·) of each subtask (including SR_Go and NR_Go). For a given subtask *t*, the similarity score between ***f***(*t*,·) and 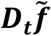 is defined as

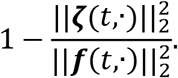

Specifically, for SR_Go and NR_Go, we separately did the approximation with the separate equation below,

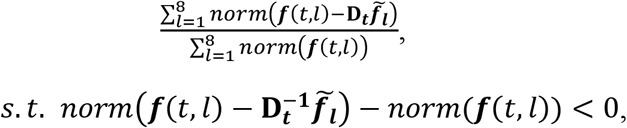

which 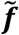 is derived from 12 subtasks approximation, *t* is for subtask SR_Go or NR_Go. Then the related similarity scores were calculated.

### Principal angles between subtask spaces

To characterize how different subtask spaces were oriented in neural state space, we computed their principal angles, a measure for quantifying the alignment of two manifolds(*64*). Specifically, given two subtask spaces *a, b* and the associated two-dimensional bases *V*_*a*_, *V*_*b*_, we performed SVD onto the inner product matrix 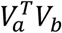, such that

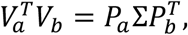

where both *P*_*a*_ and *P*_*b*_ are 2 × 2 orthogonal matrices and Σ is a 2 × 2 diagonal matrix whose elements are the ranked cosines of the principal angles *θ*_1_ and *θ* :

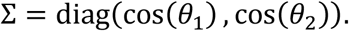

We used Σ to reflect the trend of tasks. As a control measure, we shuffled the neuron index of *V*_*b*_, ensuring that the same of projected subtask space *a*. The shuffled Σ are all below 0.05.

### Constructing a neural pseudo-population

To enable neuronal population decomposition, we constructed a synthetic population ensemble through condition-matched trial resampling. For each recorded neuron, *K* trials (resampled with replacement from trials sharing identical experimental parameters including location, subtask, and meta-task) were temporally aligned to subtask-specific anchors. This process generated a three-dimensional neural activity matrix *M*_*population*_ ∈ ℝ^*K×T×N*^, where *K* denotes condition-dependent trial count (varied across experimental conditions), *T* denotes time and *N* denotes total neuron count across recording sessions.

### Cross-task generalization on location discrimination

To test the generality of location representations across subtasks, we employed a linear support vector machine (SVM) to classify location information in each subtask. We fitted the linear-SVM with averaged neuronal activities during 330-1000 ms post-stimulus in NR_Go, SR_Go, AT_Go and DT_Go subtasks, and applying the fitted model to classify the locational information of the neuronal activities from the same averaged epochs in each subtask and the averaged baseline (100-800 ms before stimulus onset) activities in AT, DT and NR tasks. To statistically measure the generality of locational representations across subtasks, we resampled 50% of trials in NR_Go, SR_Go, AT_Go and DT_Go subtasks to construct training set *M*_*train*_ ∈ ℝ^*(×8×K)×N*^ for 100 times, where *S* denotes different subtasks. And the remaining 50% trials were used to establish test set *M*_*test*_ ∈ ℝ^*(8×K)×N*^ for each subtask in the corresponding bootstraps. For the remaining subtasks which did not participate in the fitting model, we used all trials to establish the test set for each subtask. Both training and test sets were designed as location-balanced datasets. The construction of the sets described above is detailed in the *Constructing a Neural Pseudo-Population* section. For each bootstrap, we shuffled locational labels to generate label-shuffled linear-SVM to classify the same test set as SVM decoder trained by true-label dataset. We applied permutation test for 10,000 iterations to statistically compare the decoding accuracies between SVMs fitted by the true and shuffled datasets. The p-values were corrected with bonferroni correction, where the adjusted threshold was defined as ∗ *p*_*adj*_ *= 0*.*05/k*, ∗∗ *p*_*adj*_ = 0.01/*k*,∗∗∗ *p*_*adj*_ = 0.001/*k* (*k* is the count of comparisons). All model training were conducted with the ‘scikit-learn’ library in Python(*65*).

### Neuron ablation in cross-task generalization decoding

By ranking the weights of the decoder constructed in Fig. 2f, we derived a curve that represents the contribution of each neuron (Extended Data Fig. 4c). Then we measured neuronal contribution to the encoding of generalized locational representations illustrated in Extended Data Fig. 4c by ranking the weights of pseudo-population neurons within the decoder. To probe the role of each neuron in the linear-SVM decoder, we designed a kind of neuron ablation method by sequentially shuffling neuronal activities based on their ranking. Firstly, we constructed a partially shuffled dataset by randomly reordering the trial arrangement, with task-factor labels kept unchanged for capturing the locational information from the remaining neurons, across all subtasks and locations for the top *N* neurons which made the strongest contributions to the decoder. Then, we explored the decay on decoding power with different number of neurons that were ablated following the decoding procedure in the ‘Cross-task generalization on location discrimination’ section (The model fitting was performed with partially shuffled datasets, but the test sets were still resampled from raw data with original trial arrangement). By iterating the procedure, we can derive the distribution of sequential shuffles across all neurons.

### Representational separation analysis

To measure the geometric dissimilarities between each pair of subtask spaces and the distance between each offset, we calculated an orthogonality score and the correlation distance between each pair of the 15 subtasks. The 15 subtasks are AT_Cue, AT_Go_in, AT_Go_out, DT_Go, DT_Hold, DT_Hold_N, SR_R, SR_N, NR_DRS, NR_R1, NR_R2, NR_Go, NR_R1_S, NR_R2_S and NR_Go_S. Notably, SR_Go was divided into SR_N and SR_R, for the reward predication would largely modulate location representations in SR_Go (Fig. 2b).

### Orthogonality score

To further quantify the alignment of different subtask spaces, we applied a measurement called the variance-accounted-for (VAF) ratio as follows. For any given spatial location *l*, the signal projected onto subtask *t* space was denoted as

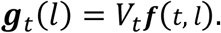

The VAF ratio with respect to subtask *a* and -*b* space-pair were defined as

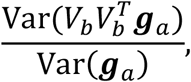

where ***g***_*a*_ = [***g***_*a*_(1), ⋯, ***g***_*a*_(8)]. Note that the VAF ratio depends on the order of *a* and *b*. Then we used 1 − *VAF ratio* to represent the orthogonality score within two subspaces. It equals to 1 if two subtask spaces were orthogonal, but it equals to 0 if they completely overlapped with each other.

### Subtask space distance

To further quantify the correlation distance between different subtask spaces, we calculated the correlation distance as subtask space distance, as follows. For any given neuronal biased response of subtask *i* and *j*, the distance between this subtask pair,

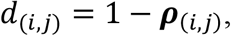

where ***ρ***_*(i,j)*_ is the Pearson correlation coefficient of between subtask *i* and *j*.

### Template-based dissimilarity regression

To assess the contributions of several potential ‘templates’ to the representation dissimilarity matrices(*34*) (RDMs), we fitted several lasso-GLM models on the vectorized orthogonality score and the correlation distance. We vectorized the design matrices of three templates: task sets, go/no-go meta-task and reward/no-reward meta-task as the factors of our model. Each of the 15*15 elements within each RSA template was explained by the following regression model:

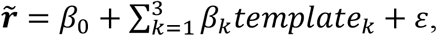

where 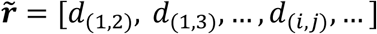 denotes the vectorized correlation-coefficient matrix from subtask space distance matrix or the matrix of orthogonality scores. Three template matrices were considered as the regressors. Then we estimated *β*_0-3_ with ordinary least squares, minimizing the sum of squared residuals, *ε*.

### Coefficients of partial determination

To quantify how these factors contributed to the encoding of subtasks, we calculated the coefficients of partial determination (CPD) for each regressor, which is defined for explanatory variable ***X***_***template***_***k***_ as follows:

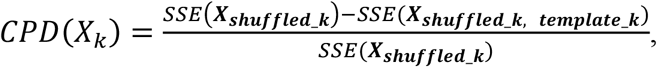

where *SSE*(*X*) refers to the sum of squared errors in the regression model that includes a set of explanatory variables ***X***_***template***_***k***_, and ***X***_***shuffled***_***k***_ is a set of all the explanatory variables included in the full model (***X***_***template***_***k***, ***shuffled***_***k***_) besides ***X***_***template***_***k***_ with shuffled trial labels.

For each subtask we had performed our linear regression model with 100 bootstrapping samples, we got the RDM and then calculated CPD values for 100 times. To statistically quantify the RDM-contribution of each template, we measured the difference between the observed and shuffled CPD values by permutation test with 1000 times of permutation on ‘observed’ and ‘shuffled’ labels.

To further detail the two monkeys’ CPD results, we separately grouped each monkey’s 100 bootstraps mentioned in “Neural state space analysis” section and then conducted the representational separation analysis on each bootstrapping sample (Extended Data Figs. 7a-7b).

For each FOV, we also applied representational separation analysis (Extended Data Figs. 7c-7f).

### Cross-task generalization on meta-task discrimination

To investigate the generality of neuronal representations about go/no-go meta-task across different subtasks, we employed SVM to classify go/no-go neuronal activities in three pairs of subtasks: 1). NR_Go belonging to go meta-task and NR_DRS belonging to no-go meta-task in the NR task, 2). AT_Go_in belonging to go meta-task and AT_Cue belonging to no-go meta-task in the AT task, and 3). DT_Go belonging to go meta-task and DT_Hold belonging to no-go meta-task in the DT task. We averaged neuronal activities during 330-1000 ms after stimulus onset for each of these six subtasks to obtain three matrices *M*_*NR*_, *M*_*AT*_, *M*_*DT*_ ∈ ℝ^*((×K)×N))*^, for three go/no-go pairs in NR, AT and DT, where 2 denotes two distinct conditions (go or no-go). We trained SVM in a ‘leave-pair-out cross-validation’ strategy: two of three pairs were chosen as the training set while the remaining pair served as the test set. Besides, an additional subtask pair (DT_Hold belonging to the reward meta-task and DT_Hold_N belonging to the no-reward meta-task) was induced as a control test group for each go/no-go discriminating decoder. Each go/no-go pair served as the test set for 100 bootstraps to establish statistical power for the estimation of go/no-go meta-task generality.

The generality of reward/no-reward was estimated following the same procedure. We chose three pairs of subtasks as the candidates for SVM training and test datasets: 1). NR_Go belonging to the reward meta-task and NR_R1 belonging to the no-reward meta-task in the NR task, 2). DT_Go belonging to the meta-task and DT_Hold_N belonging to the no-reward meta-task in the DT task, and 3). SR_R belonging to the reward meta-task and SR_N belonging to the no-reward meta-task in the SR task. The ‘leave-pair-out cross-validation’ was performed in the same manner as for go/no-go discrimination. NR_R2 subtask (go meta-task) and NR_DRS subtask (no-go meta-task) were induced as control set in each bootstrap iteration.

### Leave-one-location-out generalization test on subtasks discrimination

We selected 11 subtasks from all 14 subtasks (NR_R1_S, NR_R2_S, and SR_Go were excluded due to the insufficient trial number in the SVM training and test sets). We constructed data matrix *M*_*D*_ ∈ ℝ^*((8×11×K×N))*^ by averaging neuronal activities during 330-1000 ms after stimulus onset in 11 subtasks, where 8 denotes eight locations in each subtask. Then we selected the averaged neuronal activities of any seven locations as the training dataset *M*_*train*_ ∈ ℝ^*(7×11×K×N)*^ to fit SVM decoder for the classification of subtask information, while data of the remaining one location *M*_*test*_ ∈ ℝ^*(11×K×N)*^ was considered as test set. Neuronal activities of each individual location were utilized as test data for 100 bootstrap iterations to generate performance distributions which reflected robustness to sampling variability. Statistical significance was determined with permutation test for 10,000 iterations with bonferroni correction, where the adjusted threshold was defined as ∗ *p*_*adj*_ = 0.05/*k*, ∗∗ *p*_*adj*_ = 0.01/*k*, ∗∗∗ *p*_*adj*_ = 0.001/*k* (*k* is the count of comparisons).

### Error trial analysis in meta-task level and subtask level

For each trial, the neuronal activity was averaged from 330-1000 ms after the related event onset. To discriminate whether the neuronal responses of false-alarm trials in DT_Hold were misclassified on both meta-task and subtask levels, the linear-SVM decoders of both levels were obtained. The meta-task decoder classifying neuronal responses as go or no-go was trained on correct trials in AT_Go_in, AT_Cue, NR_Go and NR_DRS. The subtask decoder, which classified neuronal responses as DT_Go or DT_Hold was trained on correct trials in DT_Go and DT_Hold. Then the neuronal data of the false-alarm trials were applied on both meta-task decoder and subtask decoder respectively. And these data were classified as go in meta-task decoder, and as DT_Hold in subtask-decoder. As a control, the correct trials which were not included into training set were served as another test set. To measure the significance of the decoding performances, a rank-sum test was applied on each testing group.

For illustration, we first selected the data *M*_*NR*_, *M*_*AT*_ and *M*_*DT*_ described in “Cross-task generalization on meta-task discrimination” section. Then we concatenated matrices *M*_*NR*_ and *M*_*AT*_ along the first dimension, and conducted principal component analysis on the combined matrix. The first two principal components represented the space of go/no-go meta-task. Based on loadings of the PCA, the decision boundary of go/no-go meta-task decoder could be projected into the go/no-go meta-task space. Then the neuronal activity averaged across correct trials in DT_Hold and the neuronal activity averaged across false-alarm trials in DT_Hold were both projected into the meta-task spaces. Both correct and false-alarm neuronal activities during the time window from 660 ms before to 990 ms after the DT_Hold stimulus onset were selected. Then they were smoothed by a sliding window with 660 ms length and 66 ms step.

To quantify the correlation coefficients between the misclassifications of neuronal responses in the false-alarm trials of DT_Hold at both meta-task and subtask levels, we computed the distances of error trials to the decision boundaries of the decoders at both levels respectively. The distance *d* at the meta-task level could be calculated as follows:

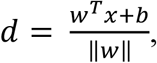

where *x* represents the neuronal responses of false-alarm trials, *w* is the weight vector of the linear-SVM decoders, *b* is the bias term, ‖*w*‖ denotes the Euclidean norm of *w*. Similarly, we calculated the distance *d*, of error trials to the decision boundaries of the decoders at the subtask level. The Pearson correlation analysis between *d* and *d*, was performed to obtain the correlation coefficients between the misclassifications on both the meta-task and subtask levels. We shuffled the indices of error trials when computing the distance from the error test data to the meta-task decoder decision boundary to obtain *d*_*shuffle*_ . The Pearson correlation analysis between *d*_*shuffle*_ and *d*′ was then performed as the control (Extended Data Figs. 8b and 8d).

To discriminate whether the neuronal responses of error trials in SR_R were misclassified on both meta-task and subtask levels, similar operations were applied to both correct and error trials in the SR_R neuronal data.

### Characterizing the functional map

To investigate whether there was anatomical organization in the encoding of task-related factors, we analyzed the presence of related functional maps across different spatial scales.

Within each FOV, we measured the anatomical organization by calculating its clustering index. First, we calculated the Pearson correlation coefficients of task-variable tuning between each neuronal pairs at location, subtask or meta-task level, respectively:

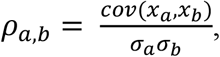

where *x* represents the vector of GLM coefficients (location level) or offsets (meta-task level and subtask level). For each meta-task type, measurements were obtained by averaging the coefficients of all corresponding subtasks for a given neuron. Both *a* and *b* are two individual neurons, and *σ* is the standard deviation of the offsets vector *x*. For a given cell-pair distance range *d* (0-50 *μm*, 100-150 *μm*, 200-250 *μm*, 300-350 *μm* or 400-450 *μm*), we quantified its clustering index by the following equation:

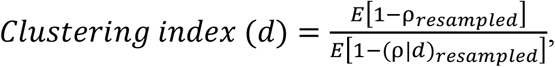

in which ρ were the correlation coefficients of all neuron pairs with all different distances, ρ|*d* were the coefficients within specific distance range *d*(*32, 66*). We randomly selected 80% of neuron pairs to generate lists of clustering indices for 100 times on meta-task level, subtask level and location level of each subtask. We also shuffled distance data across all neuron pairs to conduct a distribution of distance-shuffled clustering indices following the same protocol.

The GLM coefficients from the subtask with the best spatial codes (i.e., the largest variability of spatial locations) were used to calculate clustering index at spatial level. The GLM offsets were applied to analyzing clustering indices at subtask level. To analyze the clustering indices at meta-task level, the GLM offsets from the same meta-task group were averaged for each neuron. Then the four groups (reward group, no-reward group, go group and no-go group, the offsets of subtasks within each group were averaged as the measurements on the meta-task level) of averaged GLM offsets were used to calculate clustering indices on the meta-task level. To characterize the location clustering indices across all FOVs, the data of the subtask with the strongest spatial codes were selected.

To illustrate the anatomical organizations at subtask and meta-task level respectively (Figs. 6b-6c), the GLM offsets were normalized based on the largest absolute value within the FOVs across all bootstraps, subtasks and neurons. Then we averaged clustering indices across all bootstraps as the GLM measurements for all neurons in each subtask. For location level (Fig. 6a), the GLM coefficients were normalized for each subtask and FOV, respectively, and the averaged regression coefficients from FOV 05 on NR_Go subtask were recruited as the example.

To characterize the different levels of clustering among different levels of hierarchy in the example FOV (Fig. 6g), a permutation test was conducted between the clustering indices on location and subtask levels. Another permutation test was conducted between subtask and meta-task levels. In detail, group labels were shuffled to generate two new groups of data whose difference would be calculated between the mean values for 100,000 times, then the proportion that difference between shuffled data was greater than the difference between original groups was calculated:

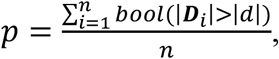

in which *n* = 100, 000, *d* is the difference of original data, ***D*** is the vector containing the differences between shuffled groups of 100,000 repetition.

To characterize the different levels of clustering among different levels across all FOVs (Fig. 6h), the data from the subtask with best spatial codes were selected. A linear model was fitted to the clustering indices as a function of hierarchical level. The averaged clustering indices across 100 bootstraps of each FOV on different hierarchies were served as *Y* -value, the group information corresponded to X-value, labelling location clustering data as 1, the subtask data were 2, the meta-task data as 3. The fitting processes were conducted by ‘regress’ function in MATLAB.

### Illustration of behavior and neural responses in movies

To illustrate a high-quality video of imaging view, we applied motion-correction and noise-reduction process on raw imaging data with CaImAn toolbox to extract denoised neuronal-activity fluorescence and background data separately. For each frame, we populated denoised neuronal-activity data into corresponding pixels in the background. And the illuminance values of each pixel were smoothed in temporal dimension by averaging the brightness values of same pixel across 1 frame before and after the current frame.

However, the contrasts of denoised video were too low to demonstrate the fluctuations of neuronal activity clearly. To enhanced video contrast, we dimmed the shadow area with nonlinear transformation:

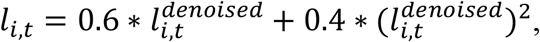

in which *l* is the brightness ranged from 0 to 1, the higher *l* value represents the brighter illuminance. *i* and *t* are given pixel and frame, respectively.

The video outputs were played with 6 frames per second (FPS), which was 0.4 × low playing-speed to raw data recording (15 FPS).

## Extended Data

**Extended Data Fig. 1.**
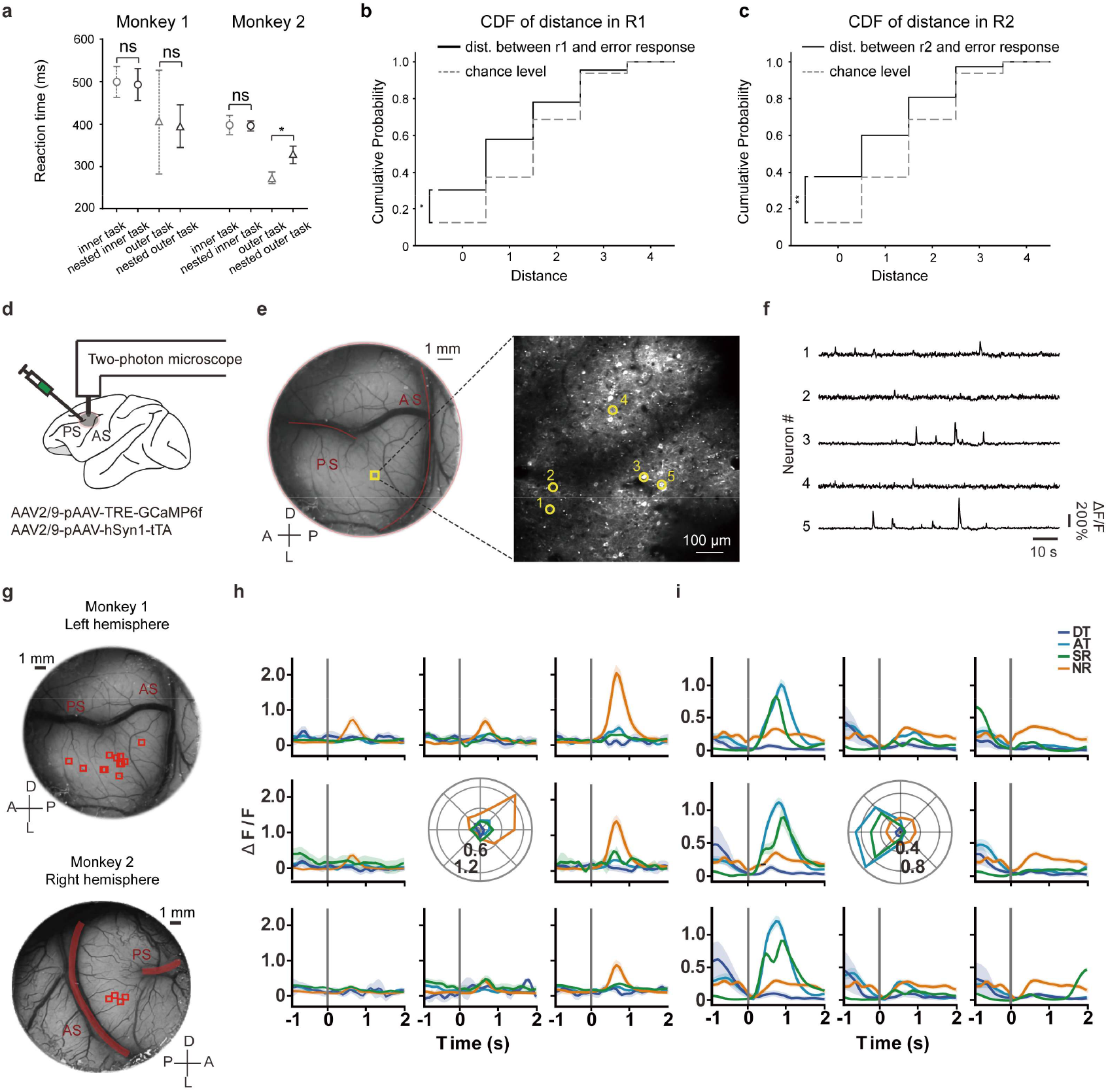
Distributions of reaction times and error responses in nested outer task in monkey. Although Monkey 2’s incorrect responses during the nested-reach go epoch (NR_Go) were not significantly aligned with the inner task targets (R1/R2) within the same trial (Fig. 1f), they exhibited a tendency to cluster around the correct targets, suggesting subtle interference from the inner task on the outer task. To further investigate this, we calculated the distances between incorrect delay responses and R1/R2 and compared their cumulative distributions to chance. **(a)** RTs across four conditions, averaged across imaging sessions. Error bars indicate SD. A significant difference was observed only between the normal outer and nested outer tasks in Monkey 2 (p = 0.029, Wilcoxon signed-rank test). **(b)** Proximity of incorrect responses to R1 target in monkey 2. Cumulative distribution of distances between incorrect delay responses and the R1 target location. The black curve represents the observed distribution, while the gray curve shows the distribution expected by chance. A significant difference was found between the two distributions (*p* = 0.016, Kolmogorov-Smirnov test). **(c)** R2 response in monkey 2. Same analysis as in **(b)**, with a significant difference between the distributions (p = 0.0015, Kolmogorov-Smirnov test). **(d)** Illustration of two-photon calcium imaging of the monkey LPFCs. AS, arcuate sulcus; PS, principal sulcus. **(e)** An example FOV, highlighted by the yellow square (left), with an enlargement (right). D, dorsal; L, lateral; A, anterior; P, posterior. **(f)** Normalized calcium traces of five example neurons (yellow circles in **(e)**), showing ΔF/F (normalized fluorescent intensity). **(g)** Distribution of FOVs in monkey 1 (top panel) and monkey 2 (bottom panel), marked by red squares. PS and AS are indicated by red lines. D, dorsal; L, lateral; A, anterior; P, posterior. **(h)** A mixed-selective example neuron with location selectivity only in NR. **(i)** An example neuron whose spatial tuning is modulated by task context, exhibiting both scaling changes and shifts in location preference across tasks.

**Extended Data Fig. 2.**
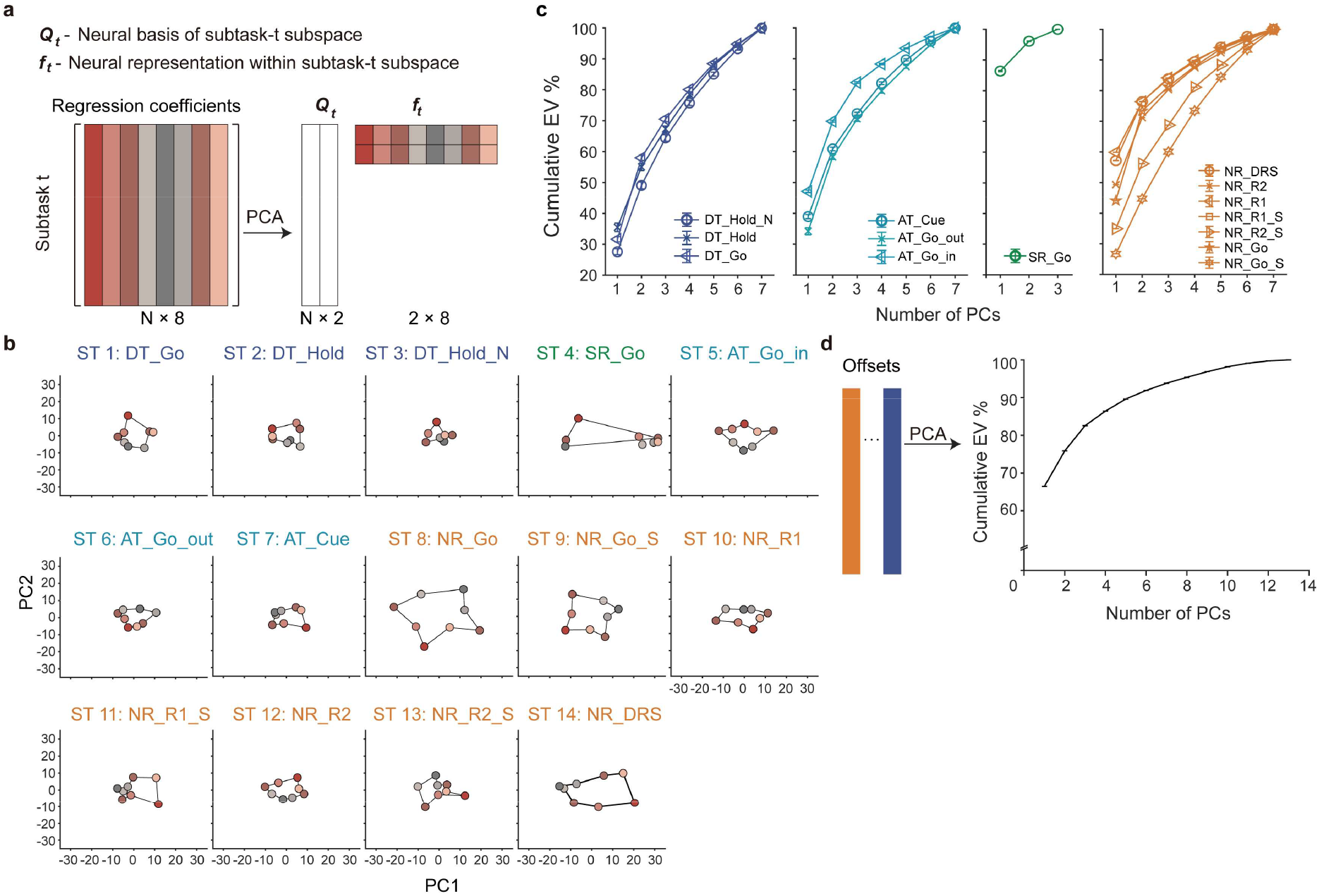
Geometric representation of multiple tasks with TDR process. **(a)** Illustration of targeted projection of neural data onto a two-dimensional subtask-*t* space using PCA analysis. **(b)** Population responses for a given task-location combination, projected onto the corresponding subtask space. Neural responses were grouped post-stimulus onset for each subtask, with locations color-coded. Axis labels in each panel correspond to the principal components related to specific subtasks. **(c)** Cumulative explained variance for PCA applied to regression coefficients from each GLM. Colors represent different tasks. **(d)** Cumulative explained variance for PCA performed on offsets from all subtasks.

**Extended Data Fig. 3.**
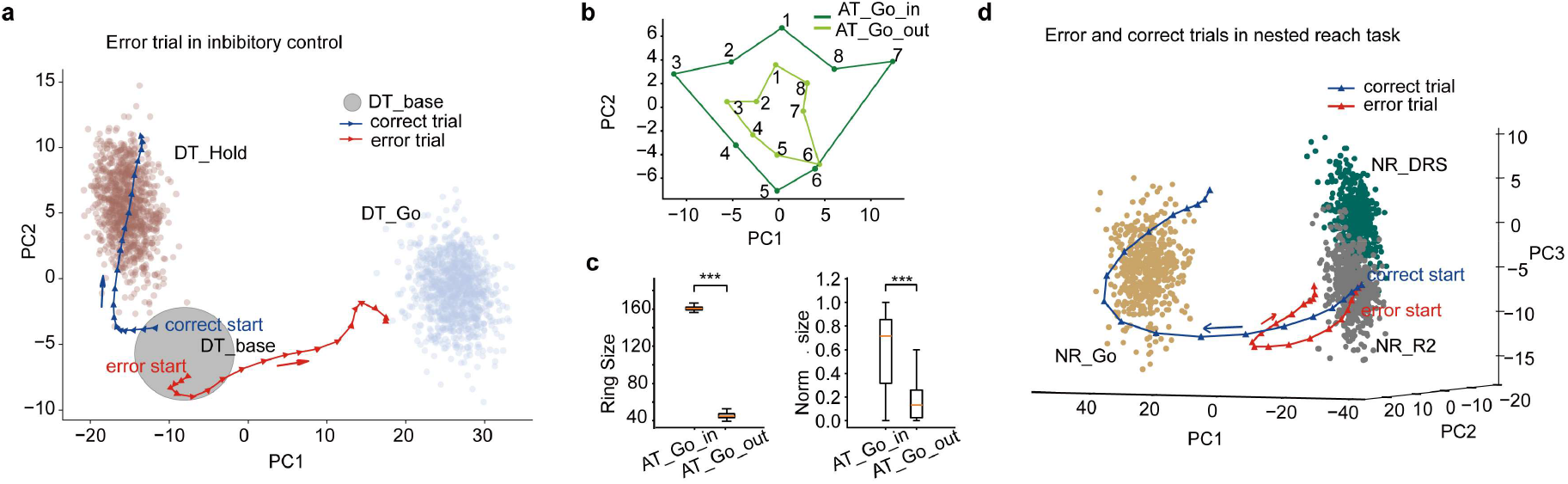
The neural geometry showed good behavior relevance. **(a)** The averaged neuronal trajectory for false-alarm trials deviates from that of correct-rejection trials. PCs represent the first two components derived from PCA on the offset terms of DT_Hold, DT_Go, and DT_base (see Methods). Projected data for different subtasks are color-coded, with each dot representing a single trial. Colored lines indicate averaged neural responses for false-alarm (red) and correct-rejection (blue) trials, and arrows denote time progression. **(b)** Comparison of geometric representations of spatial locations at the population level across different attention conditions. Neuronal responses from attend-in trials (from AT_Go_in) and attend-out trials (from AT_Go_out) are projected into a shared 2D space (see Methods). **(c)** Comparison of geometric size within the shared space between attend-in and attend-out trials. Left: Comparison based on the full pseudo-population, with size measured by projections within the space estimated from each bootstrapping sample. Geometric sizes for attend-in trials were significantly larger than for attend-out trials (160.73 ± 2.18 vs. 45.54 ± 3.31, p<0.001, t-test). Right: Comparison based on individual sessions, with size measured using the space extracted from neurons collected in each FOV, followed by normalization of bootstrap values for both subtasks within each FOV. Normalized geometric sizes for attend-in trials were significantly larger than for attend-out trials (0.60 ± 0.30 vs. 0.17 ± 0.17, p<0.001, t-test). For all panels, error bars indicate *std*, *p < 0.05, **p < 0.01, ***p < 0.001, and *ns* indicates not significant. **(d)** Comparison of neural trajectories between correct and error trials in the nested reach task. The first three PCs capture the low-dimensional manifold explaining the most variance of subtasks in the nested reach task (Methods). Projected data for different subtasks are color-coded, with each dot representing a single trial. Colored lines indicate averaged neural responses for correct (blue) and error (red) trials from the onset of DRS_Go, with arrows indicating time progression.

**Extended Data Fig. 4.**
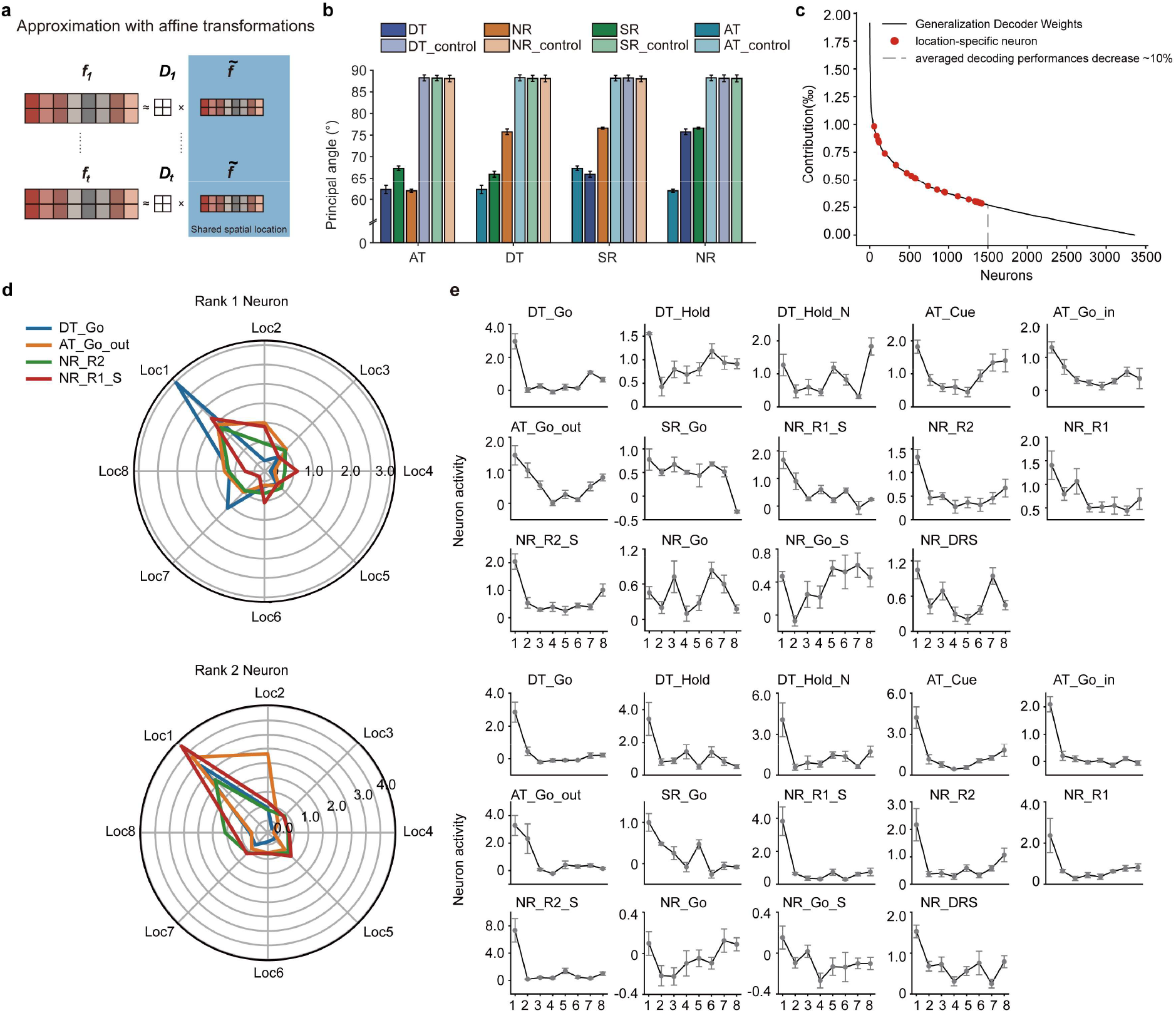
Representational sharing across multiple subtasks. **(a)** The neural representation of spatial locations within each subtask space is viewed as the outcome of an affine transformation applied to a shared representational geometry of eight locations. ***f***_***t***_, neural representations of spatial locations within subtask-*t* space. 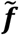, shared representational geometry across different subtasks. ***D***_***t***_, affine transformation includes both rotation and scaling. Colors represent different spatial locations. **(b)** The first principal angles between subtask spaces are shown. Tasks are color-coded, and results for control groups were obtained through random projections on each subtask space (Methods). Error bars represent standard deviation across bootstraps (n = 100). **(c)** Ranked contributions of individual neurons to the cross-task generalization decoder. Weights from the trained linear SVM decoder indicate each neuron’s contribution. The gray dashed line marks the point at which shuffling the top 1,500 neurons results in a 10% decrease in decoding performance. Red dots highlight location-specific neurons. **(d)** Spatial tuning of two top-contributing neurons in the cross-subtask generalization. Top: The first-ranked neuron shows consistent spatial preference, though with varying scales. Bottom: The second-ranked neuron exhibits a shift in spatial tuning. **(e)** Tuning curves for the two neurons with the highest contributions to the decoder across all subtasks. Each panel shows the neuron’s average activity (*mean* ± *SEM*) at each location, calculated from 330-1000 ms after stimulus onset. The top panel corresponds to the highest contributing neuron, while the bottom panel corresponds to the second-highest contributing neuron.

**Extended Data Fig. 5.**
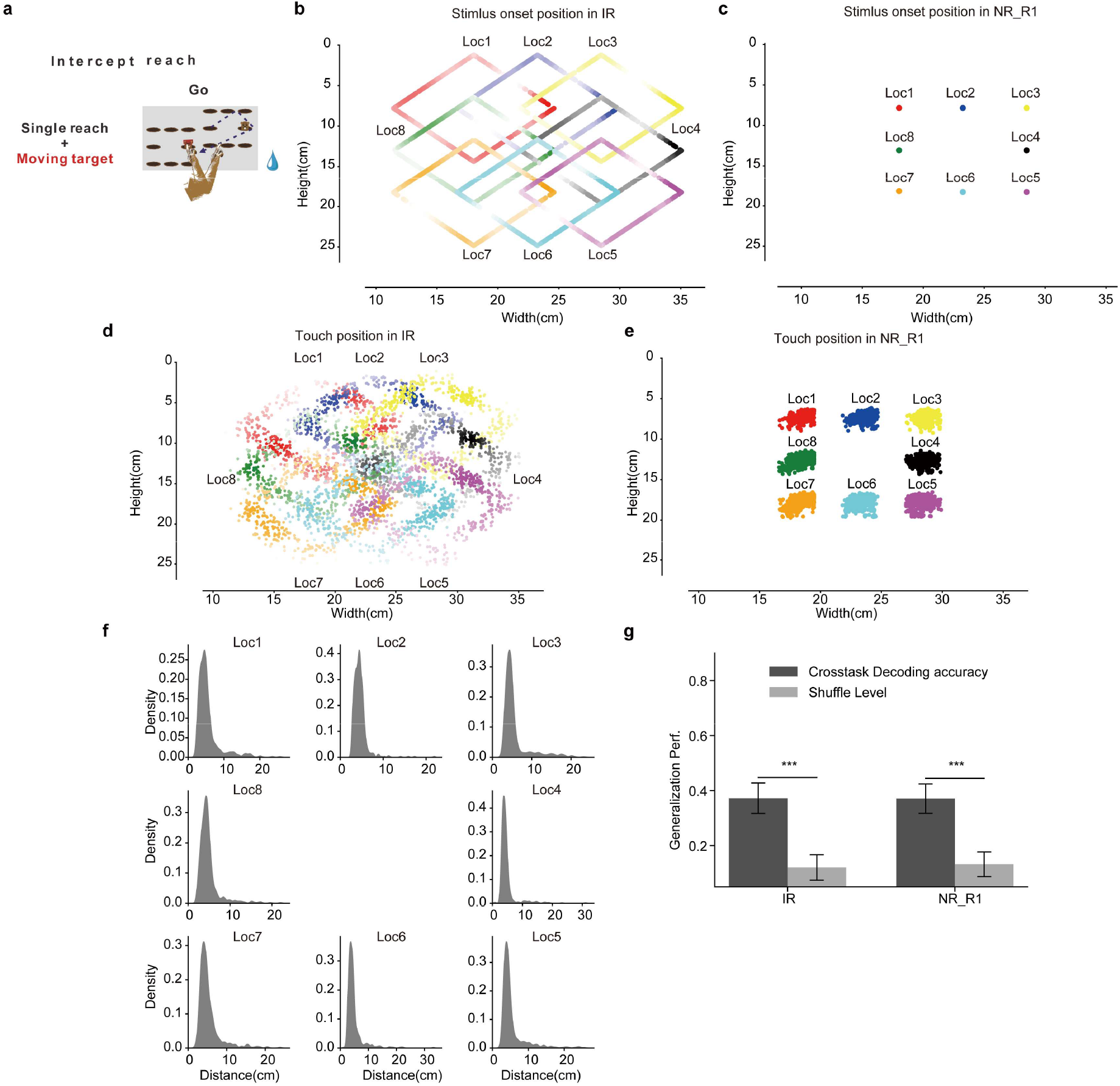
The intercept reach requires different movement patterns but with significant cross-task location decoding performance. **(a)** Task design of the intercept reach (IR) task. Both the spatial layout and the target (a mole image, i.e., the go stimulus) moved continuously along a diamond path. In each trial, both monkeys were required to touch the target within 670-4000 ms after the target appeared. For 13 sessions, the target appeared within the 670-1330 ms window, and for 3 sessions, within 2000-4000 ms window. **(b)** Density map of target positions in IR. Only the position where the stimulus first appeared was considered for each trial. The color indicates the target location, with the x-axis indicating horizontal coordinates and the y-axis indicating vertical coordinates. **(c)** Density map of target positions in NR_R1. Same analysis and format with **(b). (d)** Density map of response positions in IR. Same analysis and format with **(b). (e)** Density map of response positions in NR_R1. Same analysis and format with **(b). (f)** Distribution of distance from target onset position to touch position in the same trial. Subpanels indicate different spatial locations relative to the center of the spatial layout. **(g)** Generalization test on each subtask. Same decoder with Figure 2f was applied to the IR and NR_R1 subtasks. Both subtasks showed significantly higher accuracy than the shuffle level. *p < 0.00294, **p < 0.00058, ***p < 0.00005 (permutation test with Bonferroni correction).

**Extended Data Fig. 6.**
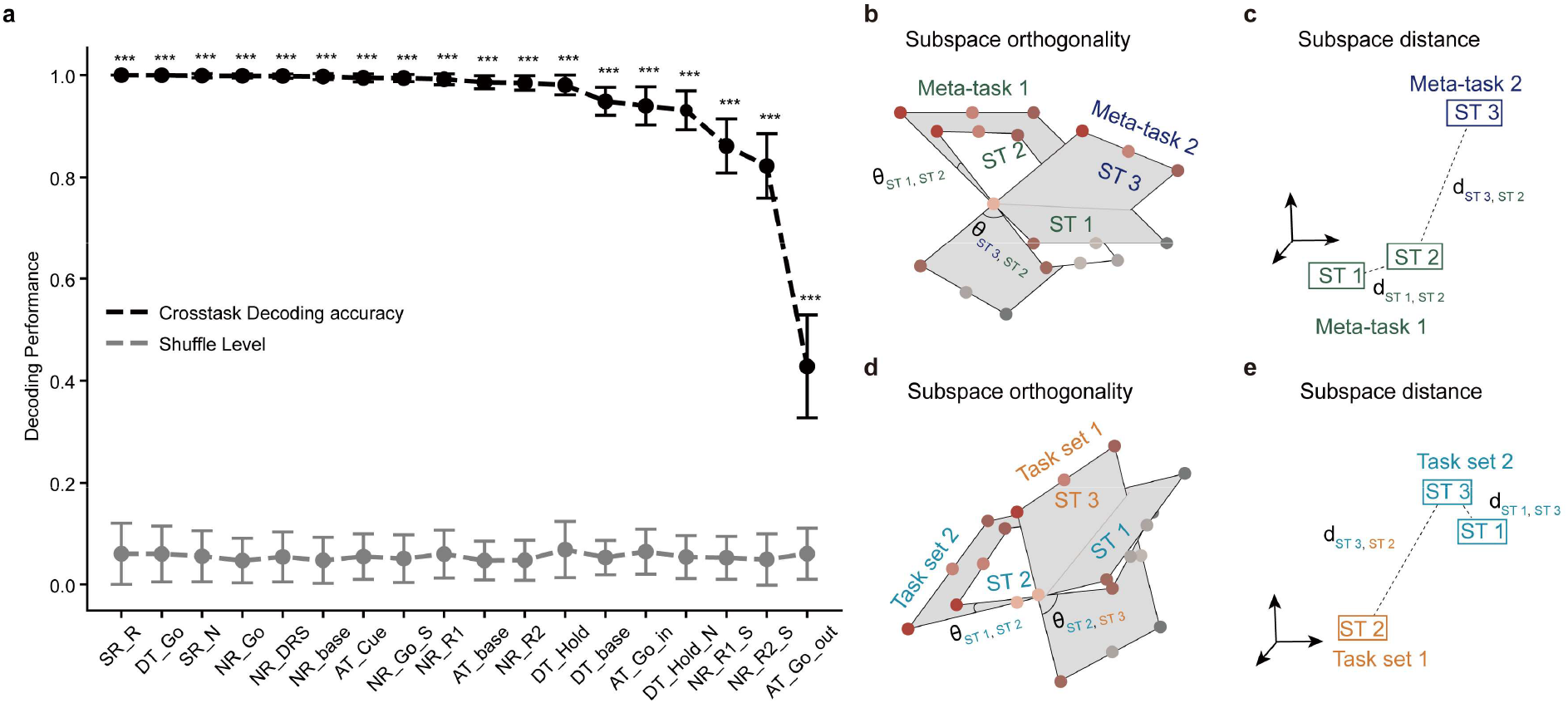
Results of subtask classification. To study the representational separation of multiple tasks, we evaluated whether neural responses are clustered within each subtask and whether subtasks are separable from one another. **(a)** Classification accuracies for 18 subtasks. Each linear SVM classifier was trained on 50% of the trials and tested on the remaining 50%. The error bars (s.t.ds) of decoding performances were estimated by 100 bootstrapping samples. Performance was compared to the shuffle level, which was established by training classifiers on shuffled labels. All subtasks were classified significantly better than the shuffle levels (Wilcoxon signed-rank test with Bonferroni level). **(b-c)** Schematic illustration of meta-tasks modulating subtask space dissimilarities. **(b)** The orthogonality score between subtasks assigned to the same meta-task is lower compared to the orthogonality score between subtasks assigned to different meta-tasks. **(c)** The subtask space distance between subtasks assigned to the same meta-task is smaller than that between subtasks assigned to different meta-tasks. Font colors indicate distinct meta-tasks, and dot colors indicate locations. **(d-e)** Schematic illustration of task sets modulating subtask space dissimilarities. **(d)** The orthogonality score between subtasks assigned to the same task set is lower compared to the score between subtasks from different task sets. **(e)** The subtask space distance between subtasks assigned to the same task set is smaller than that between subtasks from different task sets. Font colors indicate task sets, and dot colors indicate locations.

**Extended Data Fig. 7.**
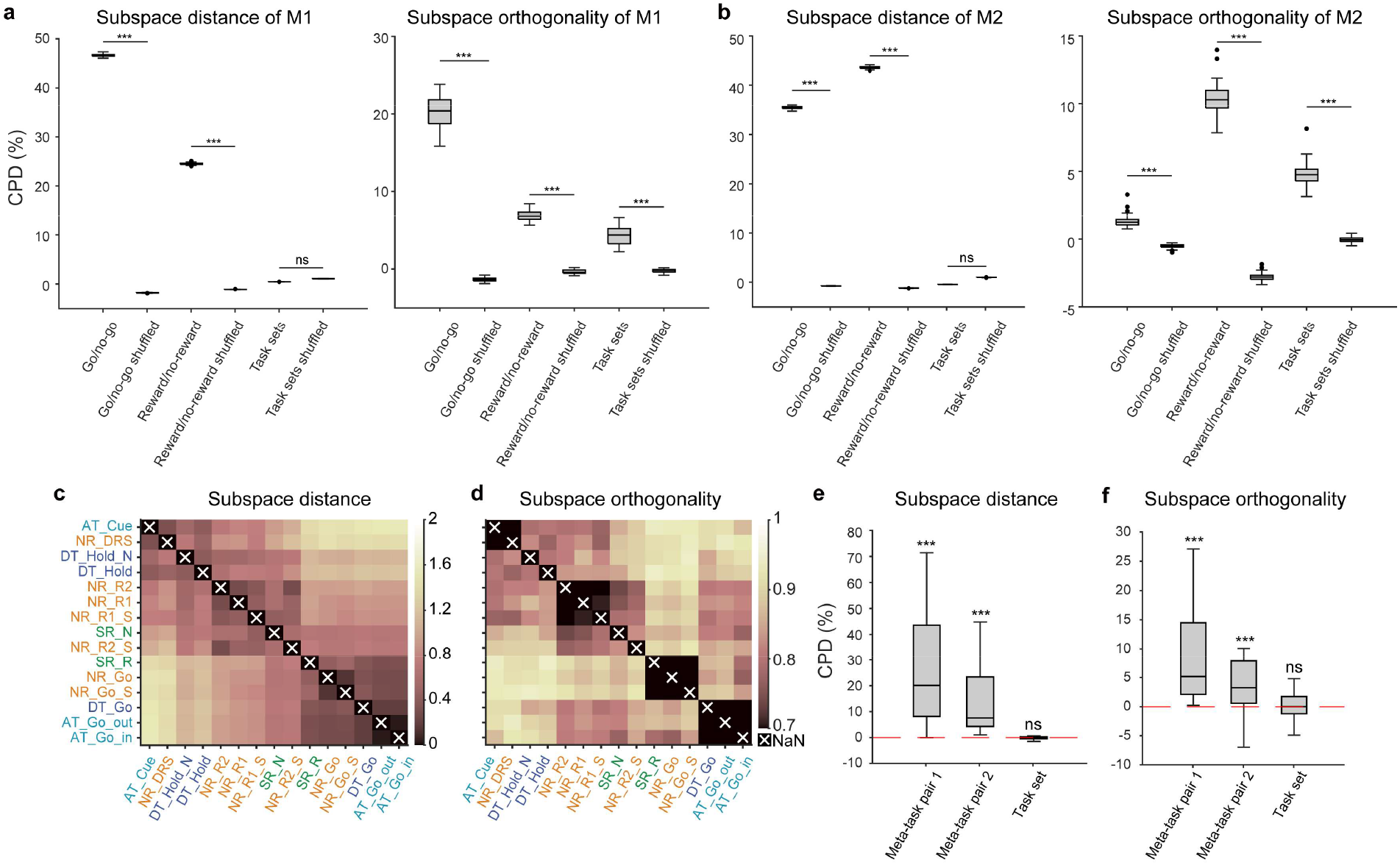
Results of representational separation analysis based on individual monkeys and individual session. To further detail the two monkeys’ CPD results, we separately grouped each monkey’s 100 bootstrapping regression results (as described in the “State space analysis” section), and then performed representational separation analysis on each bootstrapped sample. Dots indicate individual bootstrapping samples. **(a-b)** CPDs of meta-task and task set factors of monkey 1 (**a**) and monkey 2 (**b**). For both go/no-go and reward/no-reward meta-tasks, the CPDs are significantly different from 0 (*** p < 0.001), while for the task set factors, no significant difference is observed (p > 0.05). **(c)** Dissimilarities of subtask space distance. The dissimilarities of subtask space orthogonality were calculated within each session and then averaged across sessions, Same format to Figure 3b. **(d)** Dissimilarities of subtask space orthogonality. The dissimilarities in subtask space orthogonality were first calculated within each session, followed by averaging the results across sessions, same format to Figure 3a. **(e)** CPDs of meta-task and task set forneural dissimilarities of subtask space distance. The data were grouped by session and compared to the chance level. A permutation test revealed that for meta-task pair 1 and meta-task pair 2, the CPDs were significantly different from 0 (p < 0.001), while no significant difference was found for the task set (p > 0.05). **(f)** CPDs of meta-task and task set for neural dissimilarities of subtask space orthogonality. The data were grouped by session and compared to the chance level. A permutation test showed that the CPDs for both meta-task pair 1 and meta-task pair 2 were significantly different from 0 (p < 0.001), whereas no significant difference was observed for the task set (p > 0.05).

**Extended Data Fig. 8.**
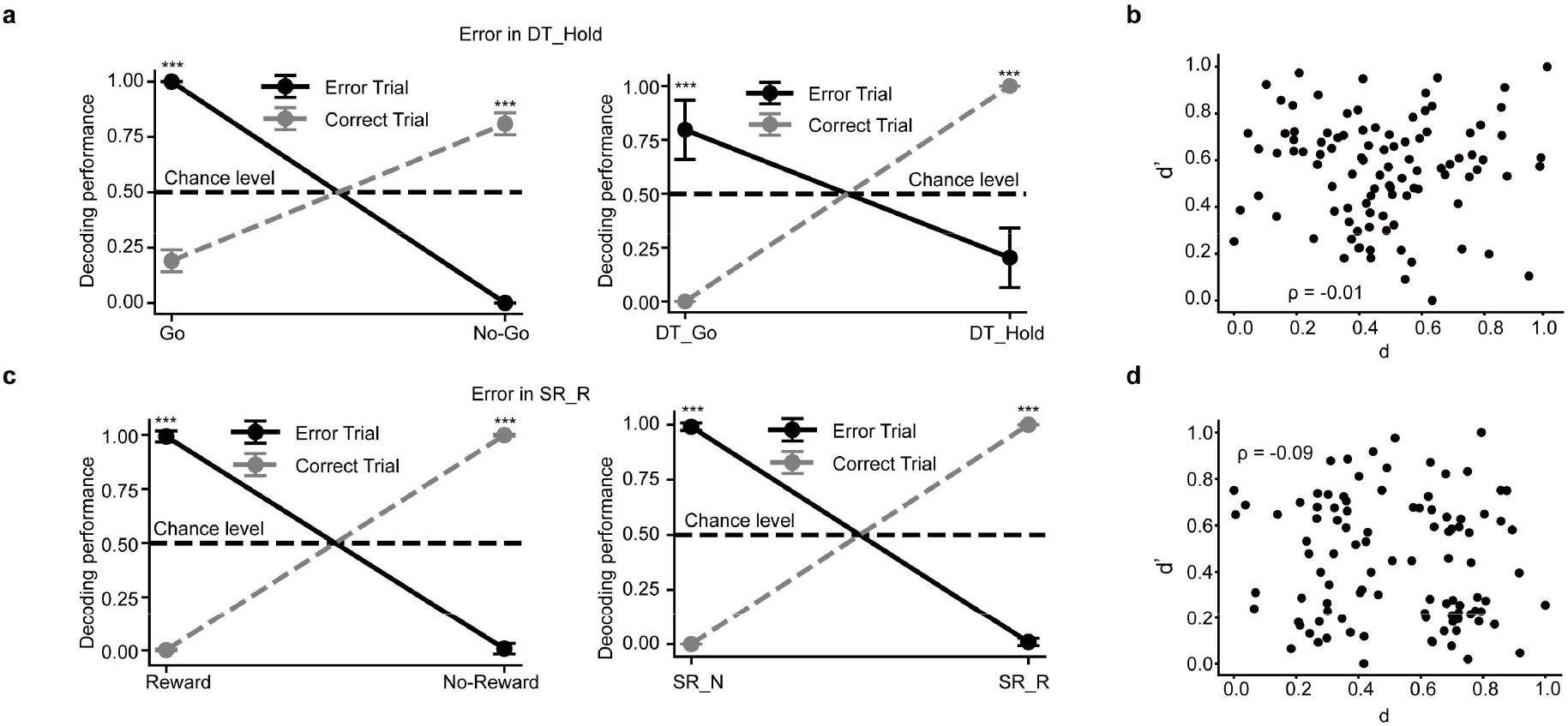
Errors occurred at both meta-task level and subtask level. **(a)** Neuronal responses of false-alarm trials in DT_Hold misclassified on both meta-task and subtask levels. Left: Decoding performance of the meta-task classifier (go vs. no-go). The accuracy of false-alarm trials classified as the go meta-task was significantly above chance (1.0 ± 0 vs. 0.5 ± 0, p < 0.001, Wilcoxon signed-rank test). Likewise, the accuracy of correct trials classified as the no-go meta-task was also significantly above chance (0.81 ± 0.05 vs. 0.5 ± 0, p < 0.001, Wilcoxon signed-rank test). Right: Decoding performance of the subtask classifier (DT_Go vs. DT_Hold). The accuracy of false-alarm trials classified as DT_Go was significantly above chance (0.80 ± 0.14 vs. 0.5 ± 0, p < 0.001, Wilcoxon signed-rank test). Likewise, the accuracy of correct trials classified as DT_Hold was significantly above chance (1.0 ± 0 vs. 0.5 ± 0, p < 0.001, Wilcoxon signed-rank test). Error bars indicate *std*, *p < 0.05, **p < 0.01, ***p < 0.001, and ns indicates no significant difference. **(b)** Scatter plot of shuffled distance pairs for error trials in DT_Hold, with a Pearson’s correlation coefficient of -0.01 (Methods). **(c)** Similar to (**a**), but with errors occurring at the reward/no-reward meta-task level and the SR_R/SR_N subtask level. **(d)** Scatter plot of shuffled distance pairs for error trials in SR_R, with a Pearson’s correlation coefficient of -0.09.

**Extended Data Fig. 9.**
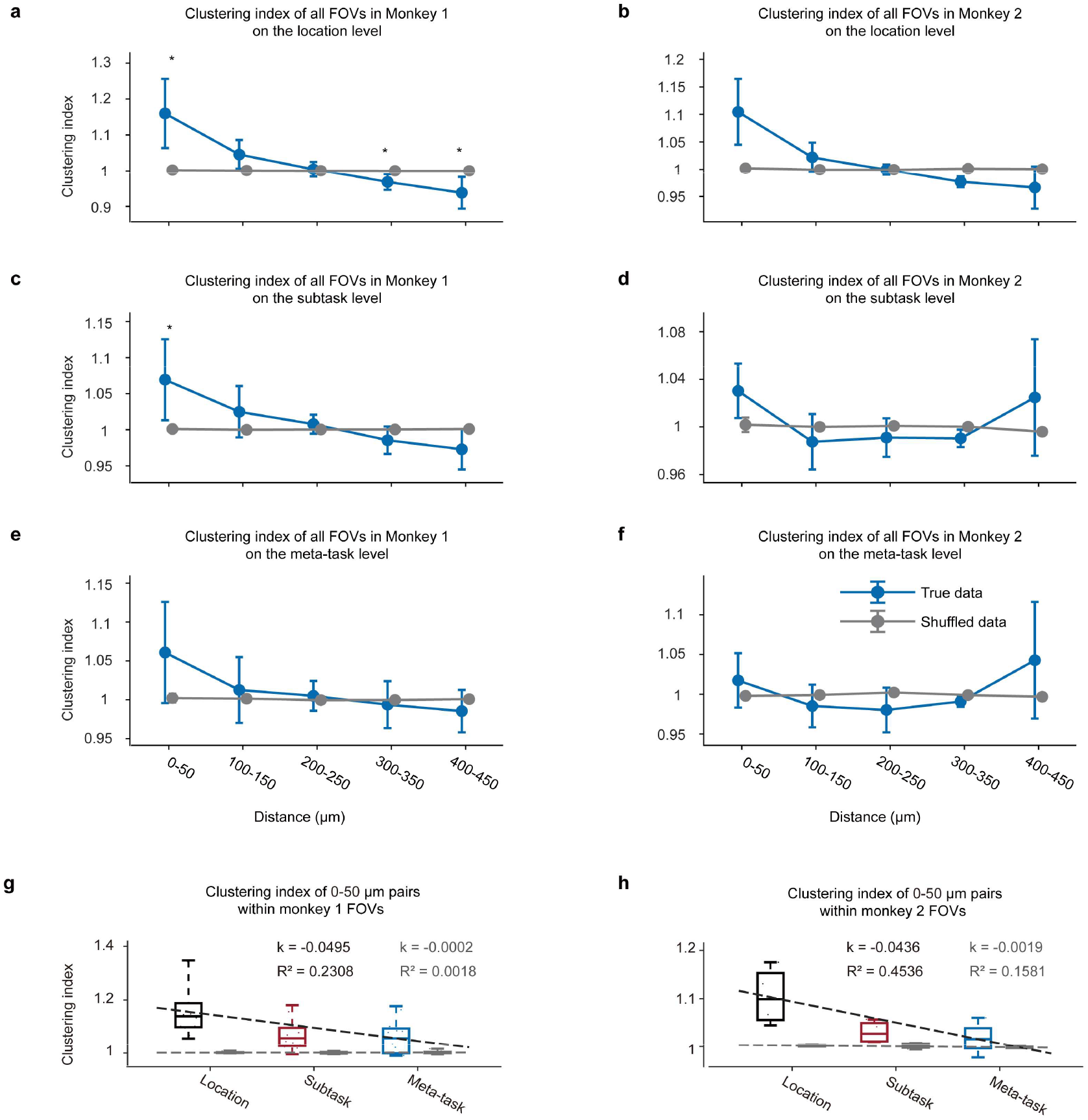
The anatomical organization on the encoding of task variables at different hierarchies. **(a-f)** Clustering index across all FOVs for both monkeys at different hierarchies (**a-b:** location level; **c-d:** subtask level; **e-f:** meta-task level; **a, c, e:** monkey 1; **b, d, f:** monkey 2). Blue line represents clustering index with true distance labels, while the gray line shows the clustering index with shuffled distance labels, permutation test, *p < 0.0001. The error bars indicate standard deviation. **(g-h)** Clustering index of 0-50 μm neuronal pairs at different hierarchies cross all FOVs of monkey 1 **(g)** and monkey 2 **(h)** at different hierarchies. The black dashed line represents the linear model fitted to the clustering index among the three groups (**g:** the slope was *k* = −0.0495 with a goodness of fit of 0.2308; **h:** the slope was *k* = −0.0436 with the goodness of fit of 0.4536). The gray dashed line represents the model fitted to shuffled data (**g:** the slope *k* = −0.0002, the goodness of fit was 0.0018; **h:** the slope *k* = −0.0019, the goodness of fit was 0.1581).

**Extended Data Table 1.** Time limit ranges for different tasks across sessions.

|  | Sessions | Response time window (ms) |  |  |  |
| --- | --- | --- | --- | --- | --- |
|  |  | Discrimination task | Attention task | Sequential reach | Nested reach (inner task) |
| Monkey 1 | 1 | 670-1330 | 330-670 | 330-670 | 670-1330 |
|  | 2 | 670-1330 | 330-670 | 330-670 | 670-1330 |
|  | 3 | 670-1330 | 330-670 | 330-670 | 670-1330 |
|  | 4 | 330-670 | 330-670 | 330-670 | 2000-4000 |
|  | 5 | 330-670 | 330-670 | 330-670 | 2000-4000 |
|  | 6 | 670-1330 | 330-670 | 330-670 | 2000-4000 |
|  | 7 | 670-1330 | 330-670 | 330-670 | 670-1330 |
|  | 8 | 670-1330 | 310-620 | 330-670 | 670-1330 |
|  | 9 | 670-1330 | 310-620 | 330-670 | 2000-4000 |
|  | 10 | 670-1330 | 330-670 | 330-670 | 500-1000 |
|  | 11 | 670-1330 | 330-670 | 330-670 | 1000-2000 |
|  | 12 | 670-1330 | 330-670 | 330-670 | 670-1330 |
| Monkey 2 | 1 | 2000-4000 | 330-670 | 330-670 | 670-1330 |
|  | 2 | 670-1330 | 330-670 | 330-670 | 1000-2000 |
|  | 3 | 670-1330 | 330-670 | 330-670 | 670-1330 |
|  | 4 | 670-1330 | 330-670 | 330-670 | 670-1330 |

**Extended Data Table 2.** Number of ROIs from each FOV.

| Monkey 1 |  | Monkey 2 |  |
| --- | --- | --- | --- |
| FOV | ROI number | FOV | ROI number |
| 01 | 255 | 01 | 269 |
| 02 | 184 | 02 | 268 |
| 03 | 191 | 03 | 236 |
| 04 | 168 | 04 | 164 |
| 05 | 203 |  |  |
| 06 | 224 |  |  |
| 07 | 199 |  |  |
| 08 | 158 |  |  |
| 09 | 200 |  |  |
| 10 | 201 |  |  |
| 11 | 244 |  |  |
| 12 | 197 |  |  |

**Extended Data Table 3.** Orthogonality scores between subtask spaces.

|  | AT<br>_Cue | NR_<br>DRS | DT_<br>Hold<br>_N | DT_<br>Hold | NR_<br>R2 | NR_<br>R1 | NR_<br>R1_S | SR_<br>N | NR_<br>_R2<br>_S | SR_<br>R | NR_<br>_Go | NR_<br>_Go<br>_S | DT_<br>_Go | AT<br>_Go<br>_out | AT<br>_Go<br>_in |
| --- | --- | --- | --- | --- | --- | --- | --- | --- | --- | --- | --- | --- | --- | --- | --- |
| **AT_Cue** | - | 0.74 | 0.85 | 0.90 | 0.85 | 0.85 | 0.84 | 0.90 | 0.95 | 0.98 | 0.98 | 0.98 | 0.98 | 0.99 | 0.96 |
| **NR_DRS** | 0.74 | - | 0.93 | 0.85 | 0.88 | 0.88 | 0.84 | 0.97 | 0.93 | 0.99 | 0.97 | 0.98 | 0.98 | 0.99 | 0.99 |
| **DT_Hold_N** | 0.85 | 0.93 | - | 0.94 | 0.91 | 0.90 | 0.94 | 0.98 | 0.96 | 0.99 | 0.99 | 0.99 | 0.98 | 0.95 | 0.97 |
| **DT_Hold** | 0.90 | 0.85 | 0.94 | - | 0.88 | 0.90 | 0.92 | 0.94 | 0.92 | 0.98 | 0.98 | 0.98 | 0.96 | 0.98 | 0.98 |
| **NR_R2** | 0.85 | 0.88 | 0.91 | 0.88 | - | 0.46 | 0.76 | 0.86 | 0.75 | 0.96 | 0.94 | 0.98 | 0.86 | 0.87 | 0.90 |
| **NR_R1** | 0.85 | 0.88 | 0.90 | 0.90 | 0.46 | - | 0.81 | 0.86 | 0.75 | 0.98 | 0.95 | 0.98 | 0.93 | 0.94 | 0.92 |
| **NR_R1_S** | 0.84 | 0.84 | 0.94 | 0.92 | 0.76 | 0.81 | - | 0.87 | 0.90 | 0.98 | 0.95 | 0.98 | 0.81 | 0.89 | 0.90 |
| **SR_N** | 0.90 | 0.97 | 0.98 | 0.94 | 0.86 | 0.86 | 0.87 | - | 0.95 | 0.91 | 0.98 | 0.97 | 0.91 | 0.91 | 0.88 |
| **NR_R2_S** | 0.95 | 0.93 | 0.96 | 0.92 | 0.75 | 0.75 | 0.90 | 0.95 | - | 0.99 | 0.94 | 0.99 | 0.92 | 0.96 | 0.94 |
| **SR_R** | 0.98 | 0.99 | 0.99 | 0.98 | 0.96 | 0.98 | 0.98 | 0.91 | 0.99 | - | 0.99 | 0.99 | 0.96 | 0.97 | 0.96 |
| **NR_Go** | 0.98 | 0.97 | 0.99 | 0.98 | 0.94 | 0.95 | 0.95 | 0.98 | 0.94 | 0.99 | - | 0.58 | 0.92 | 0.93 | 0.84 |
| **NR_Go_S** | 0.98 | 0.98 | 0.99 | 0.98 | 0.98 | 0.98 | 0.98 | 0.97 | 0.99 | 0.99 | 0.58 | - | 0.96 | 0.95 | 0.90 |
| **DT_Go** | 0.98 | 0.98 | 0.98 | 0.96 | 0.86 | 0.93 | 0.81 | 0.91 | 0.92 | 0.96 | 0.92 | 0.96 | - | 0.72 | 0.75 |
| **AT_Go_out** | 0.99 | 0.99 | 0.95 | 0.98 | 0.87 | 0.94 | 0.89 | 0.91 | 0.96 | 0.97 | 0.93 | 0.95 | 0.72 | - | 0.70 |
| **AT_Go_in** | 0.96 | 0.99 | 0.97 | 0.98 | 0.90 | 0.92 | 0.90 | 0.88 | 0.94 | 0.96 | 0.84 | 0.90 | 0.75 | 0.70 | - |

**Extended Data Table 4.** Distance between subtask spaces.

|  | AT<br>_Cue | NR_<br>DRS | DT_<br>Hold<br>_N | DT_<br>Hold | NR_<br>R2 | NR_<br>R1 | NR_<br>R1_S | SR_<br>N | NR_<br>R2<br>_S | SR_<br>R | NR_<br>_Go | NR_<br>_Go<br>_S | DT_<br>_Go | AT<br>_Go<br>_out | AT<br>_Go<br>_in |
| --- | --- | --- | --- | --- | --- | --- | --- | --- | --- | --- | --- | --- | --- | --- | --- |
| AT_Cue | - | 0.27 | 0.50 | 0.41 | 0.65 | 0.74 | 0.72 | 0.88 | 1.10 | 1.56 | 1.59 | 1.57 | 1.64 | 1.60 | 1.62 |
| NR_DRS | 0.27 | - | 0.69 | 0.52 | 0.76 | 0.84 | 0.70 | 0.98 | 1.22 | 1.43 | 1.57 | 1.55 | 1.53 | 1.49 | 1.50 |
| DT_Hold_N | 0.50 | 0.69 | - | 0.30 | 0.55 | 0.53 | 0.63 | 0.70 | 0.78 | 1.16 | 1.12 | 1.11 | 1.15 | 1.09 | 1.14 |
| DT_Hold | 0.41 | 0.52 | 0.30 | - | 0.59 | 0.66 | 0.71 | 0.66 | 0.84 | 1.21 | 1.23 | 1.22 | 1.26 | 1.22 | 1.26 |
| NR_R2 | 0.65 | 0.76 | 0.55 | 0.59 | - | 0.27 | 0.35 | 0.60 | 0.31 | 1.05 | 1.04 | 1.08 | 1.07 | 0.98 | 1.00 |
| NR_R1 | 0.74 | 0.84 | 0.53 | 0.66 | 0.27 | - | 0.46 | 0.59 | 0.48 | 1.01 | 0.96 | 0.97 | 0.97 | 0.94 | 0.94 |
| NR_R1_S | 0.72 | 0.70 | 0.63 | 0.71 | 0.35 | 0.46 | - | 0.44 | 0.48 | 1.00 | 0.86 | 0.85 | 0.91 | 0.85 | 0.86 |
| SR_N | 0.88 | 0.98 | 0.70 | 0.66 | 0.60 | 0.59 | 0.44 | - | 0.55 | 0.74 | 0.66 | 0.68 | 0.74 | 0.69 | 0.69 |
| NR_R2_S | 1.10 | 1.22 | 0.78 | 0.84 | 0.31 | 0.48 | 0.48 | 0.55 | - | 0.69 | 0.60 | 0.61 | 0.62 | 0.58 | 0.59 |
| SR_R | 1.56 | 1.43 | 1.16 | 1.21 | 1.05 | 1.01 | 1.00 | 0.74 | 0.69 | - | 0.33 | 0.39 | 0.26 | 0.24 | 0.24 |
| NR_Go | 1.59 | 1.57 | 1.12 | 1.23 | 1.04 | 0.96 | 0.86 | 0.66 | 0.60 | 0.33 | - | 0.09 | 0.25 | 0.27 | 0.27 |
| NR_Go_S | 1.57 | 1.55 | 1.11 | 1.22 | 1.08 | 0.97 | 0.85 | 0.68 | 0.61 | 0.39 | 0.09 | - | 0.27 | 0.27 | 0.25 |
| DT_Go | 1.64 | 1.53 | 1.15 | 1.26 | 1.07 | 0.97 | 0.91 | 0.74 | 0.62 | 0.26 | 0.25 | 0.27 | - | 0.15 | 0.13 |
| AT_Go_out | 1.60 | 1.49 | 1.09 | 1.22 | 0.98 | 0.94 | 0.85 | 0.69 | 0.58 | 0.24 | 0.27 | 0.27 | 0.15 | - | 0.02 |
| AT_Go_in | 1.62 | 1.50 | 1.14 | 1.26 | 1.00 | 0.94 | 0.86 | 0.69 | 0.59 | 0.24 | 0.27 | 0.25 | 0.13 | 0.02 | - |

## Notes

### Competing Interest Statement

The authors have declared no competing interest.

### Summary of Updates

Figure 2e revised: x tick labels sorting changed, the 3rd label is to be DT_Go, and the 4th label is to be AT_Go_out.

